# Benchmarking sample pooling for epigenomics of natural populations

**DOI:** 10.1101/2023.11.23.568483

**Authors:** Ryan J. Daniels, Britta S. Meyer, Marco Giulio, Silvia G. Signorini, Nicoletta Riccardi, Camilla Della Torre, Alexandra A.-T. Weber

**Author notes:** **Glossary:** β: Percent methylation estimated at a locus. Computational pooling: *Post-hoc* pooling of individual library epigenomic data using bioinformatics. Coverage/read depth: Number of reads used to inform a methylation estimate at a position/site. CpG: A dinucleotide sequence of 5ʹ–CG–3ʹ within a DNA molecule. DMR: Differentially methylated region (R), a region being a contiguous stretch of DNA. DMS: Differentially methylated site or single base pair position. DNAm: DNA methylation. Eco-epi: Ecology-epigenetic; A research field at the intersection of ecology and epigenetics. EM-seq: An enzymatic alternative to Bisulfite treatment for DNAm detection. The process uses two enzymatic reactions: 1. to protect 5mC and 5hmC from treatment and 2. unmodified cytosines are converted to uracil. Epigenomic libraries: A collection of overlapping DNA fragments that have been treated to detect DNA methylation, together representing the total DNA methylation of one (individual library) or several specimens in a single pooled library. Linkage group: All known genes on a chromosome. Locus: A section of DNA without necessarily specifying its size or function or relative position. MDM: Mean difference in methylation. PCA: Principal component analysis. Pooling: (in the context of this study) pooling of individual DNA samples ahead of library preparation and sequencing. Region: A section of DNA made up of contiguous bases, also referred to as a haplotype. Site/base: A single base-pair. WepiGS: Whole epi-Genome Sequencing; Non-targeted (random) sequencing of epigenomic data, includes methods such as WGBS and EM-seq. WGBS: Whole Genome Bisulfite Sequencing; Non-targeted (random) sequencing of epigenomic data following bisulfite treatment.

## Abstract

Interest in the role of DNA methylation (DNAm) has grown in ecological and evolutionary research of natural populations. While researchers are typically interested in comparing population-level variation, individual sequencing is the current standard. Natural populations have low effect sizes and thus need large sample sizes to detect differences. The cost of sequencing the necessary samples can be prohibitive in DNAm work. Pooling DNA before library preparation is a powerful tool to reduce costs but no recommendations exist for DNAm pooling in ecology-epigenetics research. We test if pooled and individual libraries provide similar global and region-specific DNA methylation signals in a natural system of response to pollution. We generated whole-epigenome data for two freshwater invasive molluscs (*Corbicula flumina* and *Dreissena polymorpha*) collected from a polluted and unpolluted locality, Lake Maggiore, Italy. Our results support that pooling effectively captures the same genome-wide and global treatment-level signals as individual libraries but we note that pooled libraries yielded orders of magnitude more input data and differentially-methylated regions (DMRs) detected compared with individual libraries. We estimated greatly lower power for regions from individual libraries compared with pooled libraries. The post-hoc process of computationally pooling data from individual libraries produced results comparable to pooled libraries in volumes but had discrepancies between DMRs. We discuss the possible causes for the discrepancies and put our results in the context of the benefits and drawbacks of sample pooling for epigenomics of natural populations.

## 1 Introduction

Epigenetics is the study of the stable but reversible modifications to the chromatin that do not alter the DNA sequence itself (Fallet et al., 2020; Gallego-Fabrega et al., 2015; Paro et al., 2021). These modifications can be established and removed in response to stimuli (Paro et al., 2021) and their interactions can regulate gene expression (Fallet et al., 2020). This mechanism by which gene expression can be modified has received considerable attention (Marin et al., 2018, Mounger et al., 2021, Brander et al., 2017). DNA methylation (DNAm) is the most frequently studied epigenetic modification, particularly in ecology-epigenetic (eco-epi) research focusing on comparisons between populations, in part because of how similar the workflow is to routine population genomics (Fallet et al., 2020; Lamka et al., 2022). For instance, of the available methods, whole epigenome sequencing (WepiGS) (e.g. whole-genome bisulfite sequencing (WGBS) and whole-genome enzymatic-conversion sequencing (EM-seq) offer the highest resolution available as changes can be tracked as base-pairs across the entire genome (Fallet et al., 2020; Paro et al., 202, Ziller et al., 2014).

A recent eco-epi review highlighted gaps in taxonomic and geographic sampling, and adequate replication particularly in population-level studies (Lamka et al., 2022). Effect sizes in ecological settings tend to be small, so large numbers of samples (e.g. >100 individuals per population or condition) are required to detect differences (Lea et al., 2017). Increasing sample size is not always possible in the case of rare or endangered species, but in most cases the maximum sample size is limited by budget. Indeed, the sampling effort, individual library preparation and sequencing have a strong impact on research costs. In WepiGS studies, data is typically obtained at the individual level (i.e. resulting from individual library preparation), however researchers are usually interested in population-wide signals. While the cost of sequencing has strongly decreased since its advent (Jobling et al., 2014) and is currently still decreasing, wet laboratory costs including individual library preparation remain a major obstacle for large sample sizes in many eco-epi research projects. Hence, optimizing these steps is crucial to obtaining data with the highest statistical power in a cost-effective manner.

A way to decrease costs associated with library preparation would be to pool the DNA from individual samples from the same population or condition prior to library preparation. The pooled libraries would thus represent the average signal of the individuals contained therein, with the advantage to prepare a single library. DNA sample pooling is commonly used in population genomics, where accurate population allele frequencies can be obtained from a large number of pooled samples (Ozerov et al., 2013, Konczal et al., 2014). Furthermore, pooling can also be used in transcriptomic studies, as it has been shown that pooling RNA samples and reducing coverage are effective ways to optimize costs while maintaining sufficient power in differential expression analyses (Assefa et al., 2020). So far, few studies compared the effect of sample pooling using DNA methylation data. One of them showed consistent results between individually run samples and pooled samples, where correlation coefficients were >0.98 for CpG array data (Gallego-Fabrega et al., 2015). Two further studies focusing on MALDI-TOF mass-spectrometry data from individual and pooled DNA produced strong evidence that pooled DNA samples provide reliable estimates of group DNA methylation averages and showed that the agreement holds up with a variety of pooling sizes (Docherty et al, 2009; Docherty et al., 2010). To date, however, the comparison between individual and pooled samples has not been done with WepiGS.

While pooling samples has a strong potential for increasing power and decreasing costs, there are important considerations related to methylation data which have led to recommending against sample pooling (see Ziller et al., 2014; Lea et al, 2017). First, methylation data is more variable than genomic data by virtue of its inducibility and reversibility (Tsai & Bell, 2015). Therefore, individual samples are not only snapshots in time and space, but they also represent the somatic average. Second, methylation patterns may be tissue specific (Lee et al., 2017) and more closely-related cell-types have more similar methylation signals (Blake et al., 2020, Ziller et al., 2014). Thus, biases may be introduced if inter-individual (or inter-tissue) variation cannot be accounted for (Teschendorff et al., 2017). Finally, a particular concern has been that pooling masks variation, prevents inclusion of covariates (Tsai & Bell, 2015, Ziller et al., 2014), and ultimately requires more biological replicates to account for the hidden variation (Futschik & Schlotterer, 2010). Most importantly, if samples are pooled there is no possibility of going back to the individual data, so any covariation in the data that was not expected or previously identified in the original pooling design will be masked.

As the benefits and drawbacks of sample pooling in whole-genome DNA methylation studies have not been formally compared, and there are currently no clear recommendations about the pertinence of pooling DNA for epigenomics of natural populations. To address this gap, we investigated empirically the effects of sample pooling in DNAm by using two invasive freshwater bivalves from polluted and unpolluted localities as study systems. The aims were to: 1) test whether global DNAm signals from pooled and individual libraries are equivalent, 2) compare the overlap between differentially methylated regions between polluted and unpolluted localities arising from individual and pooled datasets and 3) provide a set of recommendations about the pertinence of sample pooling for future eco-epi projects.

## 2. Methods and materials

### 2.1 Sampling and DNA extraction

Individuals of the Asian clam *Corbicula fluminea* and the zebra mussel *Dreissena polymorpha* were collected by SCUBA diving at either polluted or unpolluted localities in Lake Maggiore, Italy, and frozen at −20C upon arrival in the laboratory (Table 1). Sampling permits were not necessary as both species are invasive. DNA extractions for 40 individuals (10 DP, 10 DNP, 10 CP, 10 CNP) from foot tissue were performed using DNeasyBlood and Tissue DNA extraction kit (Qiagen Cat.no. 69504) following the manufacturer’s recommendations (Elution in 110 *μ*l elution buffer). DNA integrity was examined using agarose gel electrophoresis and DNA concentration was measured using Qubit 2.0 (Invitrogen). Two *Corbicula* extractions failed (1 CP, 1 CNP), leaving 38 DNA extractions for individual library preparation (Summary of the experimental design in Figure 1).

**Figure 1:**
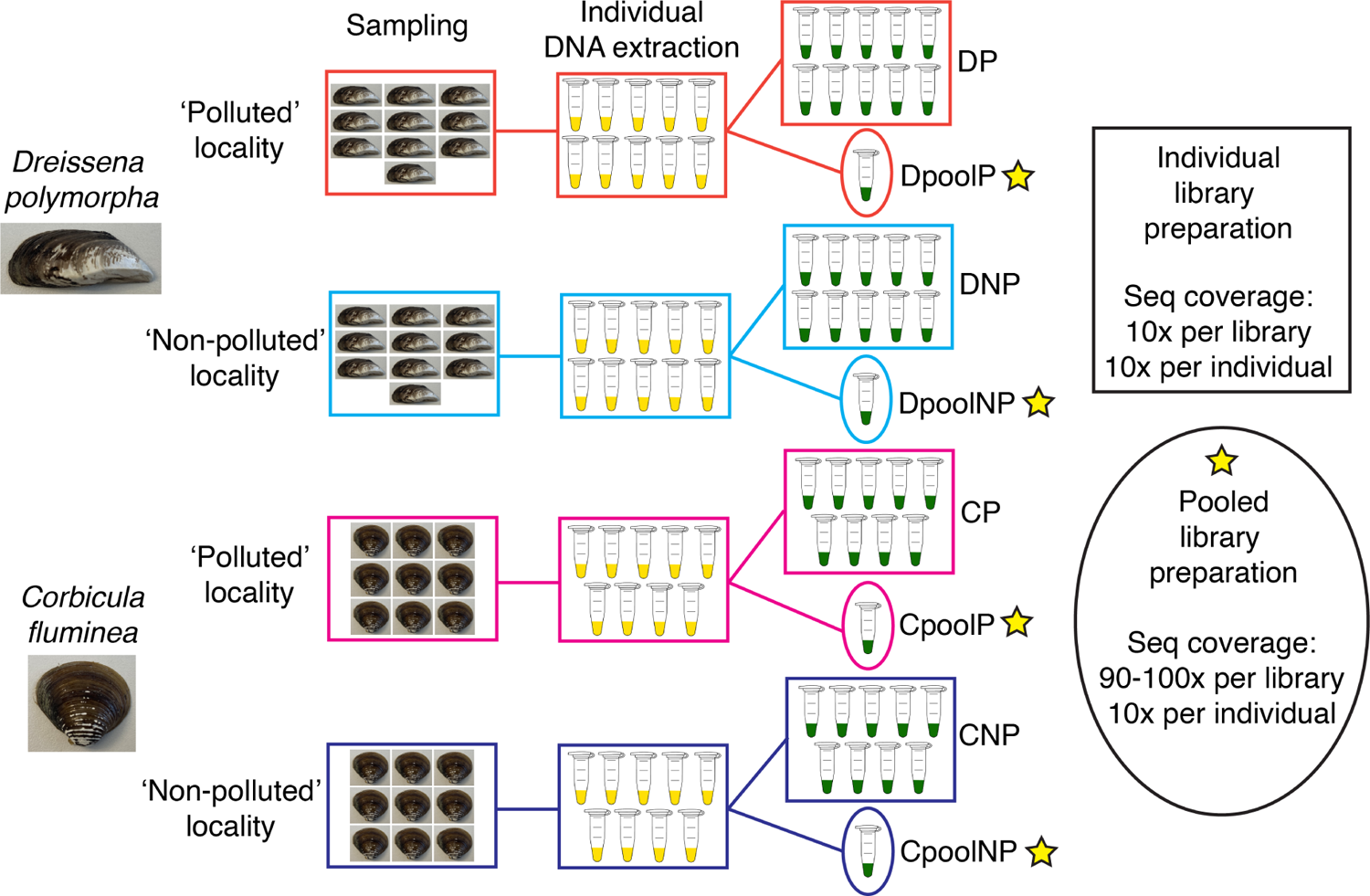
Experimental design of the study. Individuals from two species (*Dreissena polymorpha* and *Corbicula fluminea*) were collected at polluted and non-polluted localities in Lake Maggiore, Italy. See Table 1. Individual DNA extractions were performed. The same DNA extractions were used to construct 38 individual and four pooled libraries. Individual and pooled libraries were sequenced at an equivalent per-individual sequencing coverage (i.e. ∼10x per individual).

**Table 1:** Summary of sampling design. All localities are in Lake Maggiore, Italy.

| Species | No. of specimens | Locality | Coordinates<br>[decimal degrees] | Sampling date | Depth<br>[m] | Environment<br>[Treatment group] | Acronym |
| --- | --- | --- | --- | --- | --- | --- | --- |
| *Dreissena polymorpha* | 10 | Baveno | 45.915011 N, 8.503474 E | 22.05.20 | 6 | Polluted | DP |
|  | 10 | Cannobio | 46.092616 N, 8.691536 E | 22.05.20 | 6 | Non-polluted | DNP |
| *Corbicula fluminea* | 10 | Baveno | 45.915011 N, 8.503474 E | 09.05.18 | 4 | Polluted | CP |
|  | 10 | Magadino | 46.153440 N, 8.852953 E | 13.08.20 | 4 | Non-polluted | CNP |

### 2.2 Pooling design, enzymatic conversion, library preparation and sequencing

To ensure individual and pooled libraries were comparable, the pooled libraries were prepared from equimolar amounts of individually extracted DNA (Table S1; Figure 1), generating four pooled libraries. The libraries were as follows; 1) *Dreissena* pool polluted (DpoolP) representing the population of 10 *Dreissena polymorpha* individuals from the polluted locality; 2) *Dreissena* pool non-polluted (DpoolNP) representing the population of 10 *Dreissena polymorpha* individuals from the non-polluted locality; 3) *Corbicula* pool polluted (CpoolP) representing the population of 9 *Corbicula fluminea* individuals from the polluted locality; 4) *Corbicula* pool non-polluted (CpoolNP) representing the population of 9 *Corbicula fluminea* individuals from the non-polluted locality. We prepared a total of 42 libraries, 38 individual ones and 4 libraries from pooled DNA. We used an enzymatic technique to convert unmethylated cytosines in thymidine as it minimizes DNA damage. We used the NEB Next Enzymatic Methyl-seq Kit (New England Biolabs Cat.no. E7120S). Control DNA (CpG methylated pUC19 and unmethylated lambda) used to estimate conversion rates was added to each DNA extraction before shearing as per the manufacturer’s instructions (New England Biolabs). For each individual DNA extraction, DNA was sheared with a Qsonica sonicator (Q800R2 instrument) using different shearing times depending on the level of DNA integrity previously assessed using agarose gel electrophoresis: 1) samples with high-molecular weight DNA were sheared 2’45’’ minutes; 2) samples with semi-degraded DNA were sheared 9’00’’-11’30’’ minutes; 3) samples with highly-degraded DNA were not sheared. For the samples with highly-degraded DNA, control DNA was sheared individually (nine minutes) and then added to the sample DNA. Library preparation was done following the manufacturer’s instructions except that we used half volumes of all reagents to reduce costs. The reduced reagent volumes did not have any clear impact the sequencing output.

Final libraries were checked on a TapeStation 4150 Instrument (Agilent Technologies). We aimed to obtain the same mean coverage per sample from both the individual libraries and the pooled libraries of each species. We thus combined individual libraries in equimolar concentrations and the pooled libraries in a molar concentration x-fold higher than the individual libraries (i.e. 10x for *Dreissena polymorpha* pools and 9x for *Corbicula fluminea* pools) as the individual and pooled libraries of a particular species were sequenced one the same lane. The 42 libraries were sequenced on two lanes of a S4 flowcell on an Illumina Novaseq 6000 sequencer (150 bp paired-end) at the Functional Genomics Center, Zürich.

### 2.3 Quality control and mapping

In total, 18 *Corbicula fluminea* individuals and 20 *Dreissena polymorpha* individuals were sequenced at an average of 74 (*±* 9.3) million reads (Table S1). The four pooled libraries were sequenced at an average of 620 (*±* 66) million reads. The reads were quality-assessed using FastQC v.0.11.9 (Andrews, 2019) and MultiQC v.1.9 (Ewels et al., 2016). Adapters were identified and removed using Trim Galore! v.0.6.6 (Krueger, 2020) with default settings. To correct for bias of methylation percentage at the read ends, reads were trimmed of 10 bases on both the 3’ and 5’ ends (as recommended; https://felixkrueger.github.io/Bismark/bismark/library_types/). Default settings were retained for all other trimming steps, including the removal of low-quality bases (–quality 20) and dropping reads shorter than 20 bases (–length 20). Enzyme conversion efficiency was assessed using the two control DNA. The high quality reads having passed QC were then aligned to the respective publicly available reference genomes; *D. polymorpha* (McCartney et al., 2022) and *C. fluminea* (Zhang et al., 2021).

Alignment, de-duplication and methylation extraction were performed with Bismark v.0.19.0 (Krueger & Andrews, 2011). Briefly, we first converted reference genomes computationally for alignment and then indexed using Bowtie2 v.2.4.4 (Langmead & Salzberg, 2012) with default settings (command bismark_genome_preparation). Alignment was run with directionality specified using the default alignment score (–score_min L,0,-1.2). Default settings included the – exclude_overlap flag which only considers data from one of the two strands available in case of overlap between forward and reverse reads. As part of the QC for the trimmed reads, we compared the number of read-pairs, the level of read duplication and the alignment efficiency between treatment groups within species. Tests were performed using base R functions including the Shapiro-Wilk test (shapiro.test) for univariate normality (Shapiro & Wilk, 1965), the Bartlett test (bartlett.test) for homogeneity of variance (Bartlett, 1937) and the ANOVA performed using the lm and summary.aov functions.

### 2.4 Coverage filtering, estimate of data loss and computational pooling

We processed the aligned reads for CpG sites with the MethylKit R package, v.1.24.0 (Akalin et al., 2012) available through Bioconductor (Huber et al., 2015). To reduce computational load, we analyzed the first eight linkage groups from each species. We decided to retain bases with at least ten reads. We further excluded over-represented sites, which may reflect sequencing bias, by removing the sites in the 99.9th percentile of coverage. Regions of one kb size were formed as non-overlapping blocks using the tile function in MethylKit with default options (sliding windows of 1000bp and regions of 1000bp).

We performed computational pooling of individual library data to allow for a better comparison with the wet-lab pooled library data with the aggregate result of individual libraries. Computational pooling is a *post-hoc* process that sums up the coverage within each treatment using the individual library data and creates one library per treatment group or population. We used the post-QC individual libraries as input data and pooled using the pool function in MethylKit. We considered two thresholds for retaining loci; only loci with an overlap of at least 75% of the samples at each locus and only loci with 100% of the samples at each locus.

### 2.5 Evaluation of concordance between pooled and individual libraries

#### 2.5.1 Genome-wide CpG methylation levels

As a test for agreement between the pooled and individual libraries, we fit an overall correlation of the CpG methylation estimates for all samples in a pairwise fashion using Pearson’s correlations with the getCorrelation function in MethylKit (Table 2). From this we could observe any bias introduced by a particular subset of regions. In the first scenario we compared all libraries against each other. In the other two scenarios we merged only the pollution and non-pollution treatment groups, respectively.

**Table 2:** Summary of data loss throughout QC. Presented are the mean number of CpG sites before and after applying a filter for 10x minimum coverage, as well as the amount of data lost as a percentage.

| Species | Libraries | Pool | Treatment | No. before filter | S.D. | No. after filter | S.D. | % loss | S.D. |
| --- | --- | --- | --- | --- | --- | --- | --- | --- | --- |
| *C. fluminea* | 7 | No | P | 15,955,679.00 | 1,184,713.97 | 703,178.29 | 247,309.93 | 96% | 0.01 |
|  | 7 | No | NP | 16,598,487.86 | 664,380.62 | 829,556.29 | 260,670.42 | 95% | 0.01 |
|  | 14 | No | P+NP | 16,277,083.43 | 924,547.29 | 766,367.29 | 253,990.18 | 95% | 0.01 |
|  | 1 | Yes | P | 23,887,527.00 | - | 15,535,134.00 | - | 35% | - |
|  | 1 | Yes | NP | 24,073,697.00 | - | 15,626,885.00 | - | 35% | - |
|  | 2 | Yes | P+NP | 23,980,612.00 | 131,642.07 | 15,581,009.50 | 64,877.75 | 35% | 0 |
| *D. polymorpha* | 8 | No | P | 30,625,991.00 | 2,369,570.65 | 844,482.00 | 261,373.55 | 97% | 0.01 |
|  | 10 | No | NP | 32,947,578.60 | 1,105,154.88 | 1,230,564.60 | 486,459.13 | 96% | 0.01 |
|  | 18 | No | P+NP | 31,786,784.80 | 1,737,362.76 | 1,037,523.30 | 373,916.34 | 97% | 0.01 |
|  | 1 | Yes | P | 48,459,924.00 | - | 33,839,276.00 | - | 30% | - |
|  | 1 | Yes | NP | 47,975,036.00 | - | 31,541,434.00 | - | 34% | - |
|  | 2 | Yes | P+NP | 48,217,480.00 | 342,867.59 | 32,690,355.00 | 1,624,819.66 | 32% | 0.03 |

To describe the relationship between the signal in the pooled and individual libraries and between the polluted and non-polluted treatments, we performed a clustering based on the % methylation estimates using a principal component analysis (PCA). To estimate an error on the PCA coordinates, we performed a jackknife over linkage groups, estimating the standard error (Busing et al., 1999; as in Montinaro et al., 2015). To confirm that jackknife iterations were reporting a similar clustering signal, we tested for a correlation between the PC loading matrix across jackknife iterations using the Tucker’s coefficient (Lorenzo-Seva & Berge, 2006; Peres-Neto & Jackson, 2001).

To prevent any conflicts in the directional of components between jackknife iterations, we used a Procrustes transformation to align each iteration of the PCA and the PCA of the full dataset (Peres-Neto & Jackson, 2001). The transformation coefficients were examined to ensure that no matrix needed excessive transformation to align as this would indicate a big difference in the signal. Where PCs were strongly correlated across jackknife iterations, we proceeded to estimate the error.

#### 2.5.2 Differential methylation in response to pollution

We tested if there was overlap between regions showing differential methylation between polluted and non-polluted localities from the individual, pooled and computationally pooled libraries. Differential methylation for the individual libraries was estimated using a logistic regression (Cramer & Howitt, 2004). Here we only considered loci that were covered by at least 75 % of the samples in a treatment, respectively. This regression cannot be conducted with one sample per treatment group (i.e. pooled libraries), so differential methylation was estimated for the pooled and computationally pooled libraries using the Fisher’s Exact Test (Fisher, 1934). We set no minimum overlap for loci across samples in the computationally pooled data (Table S2). The P-values were corrected for multiple testing under a sliding linear model method (Wang et al., 2011) and we report the q-values. Regions were considered to have significant differential methylation (i.e. DMR) with *q* <0.01 and a mean methylation difference of at least 25%. For understanding the direction of hyper/hypo-methylation, all tests were performed with the following orders for treatments; “Pollution treatment” vs. “Non-Pollution treatment”. The number of regions in common between tests were visualized with ggupset (Ahlmann-Eltze, 2020) package in R.

### 2.6 Estimates of recovered power

To gauge the available power in our dataset, we estimated the recovered power per locus in the contrast of polluted and non-polluted populations. The power of a test is defined as the probability that it correctly rejects the null hypothesis when the alternative hypothesis is true. Firstly we identified invariant regions, which we considered to be regions with no MDM between the treatments, rather than regions showing no variation in methylation. This measure takes into account both variance within and between treatments and allows for some level of artificial variance due to errors. With the individual libraries power estimates were based on a t-test. Effect sizes were estimated as Cohen’s d, which is the expected difference in means divided by the standard deviation across all samples (Cohen, 1988). We estimated standard deviation following the example by Mansell *et al*. (2019). The mean difference in methylation (MDM) at each locus was based on that calculated in the estimation of DMRs, with α = 0.01 and the observed sample sizes per treatment group per species. We consider only loci with 100% overlap across all samples. The power values were calculated using the pwr.t.test function in the R package pwr (Champely, 2018). For the pooled and computationally pooled libraries, adjustments were needed to replicate the Fisher’s Exact test. With binomial count data, the variance is a function of the mean (Everitt & Torsten, 2010) and this allows us to estimate the standard deviation as the square root of the variance function using only the proportions *Variance* =*p*_1_ /_(_1 *− p*_1)_ *n*_1_ + *p*_2_ / _(_1 *− p*_2)_ *n*_2_

The effect size was estimated using the ES.h function which uses an arcsine transformation. The power was estimated using the pwr.2p2n.test function in the R package pwr. The pwr.2p2n.test test considers a two-proportion test with unequal sample sizes (i.e. coverage in this context) under the null hypothesis that there is no difference in the treatment means. The region-specific coverage value was used in the calculation.

### 2.7 Estimates of the necessary sampling effort for significant detection

We estimated the distribution of the necessary sampling effort to detect statistically significant differences between polluted and non-polluted populations at each locus. Sampling effort estimates were made with the pwr.t.test function for the individual libraries and with the pwr.2p.test function for the pooled and computationally pooled data. We set the power threshold to 80% (power = 0.8, n=NULL) in all cases and we assumed equal sampling effort. In this estimate the sampling effort for individual libraries is measured as the number of biological replicates (each providing a methylation estimate as a continuous number). Sampling effort for the pooled and computationally pooled data is measured as the interaction of the coverage and the number of biological replicates (count data as either methylated or unmethylated read).

### 2.8 Laboratory costs estimation

We summarized the costs per sample in a hypothetical scenario where 12 populations from two treatment groups have been sampled (Table 3). We estimated the cost of creating a ‘gross pooled’, ‘nested pooled’ and ‘individual libraries’ with 8, 4, and 1 individual(s) per library, respectively. The costs were based on quotes as of 2023 in Swiss Francs including local taxes. These costs exclude any procedures which are equivalent between the pooled and the individual libraries (such as DNA extraction, sample collection and DNA sequencing, assuming equivalent sequencing depth per individual).

**Table 3:** Cost comparison of pooled and individual libraries. Estimated costs per specimen in a scenario of sampling 12 populations for a total of 192 biological replicates and no technical replicates. Costs are provided in Swiss Francs (CHF) with local 2023 prices. Costs are likely to vary among countries based on local factors. Prices include local taxes. * Tapestation D1000 screen tape, reagents and consumables. ** Pipette tips, general reagents, gloves, tubes etc. *** 1*24 rxn kit for pooled and 2*96 rxn kit for individual libraries. Where costs are equal between pooled and individual libraries, we omitted such costs (i.e. individual DNA extraction, quality control, shearing, sequencing depth).

|  | Individual | Nested Pooled | Gross Pooled |
| --- | --- | --- | --- |
| **Pooling details** |  |  |  |
| Populations | 12 | 12 | 12 |
| Libraries | 192 | 48 | 24 |
| Samples per library | 1 | 4 | 8 |
| Total samples | 192 | 192 | 192 |
| **Material/Protocol total** |  |  |  |
| NEBNext® Enzymatic Methyl-seq Kit *** | 9,374.00 | 2,604.00 | 1,302.00 |
| General consumables** | 192.00 | 48.00 | 24.00 |
| Library quantification* | 192.00 | 48.00 | 24.00 |
| Library quality control* | 960.00 | 240.00 | 120.00 |
| Total | 10,718.00 | 2,940.00 | 1,470.00 |
|  | 55.82 | 15.31 | 7.66 |
| Cost ratio (Individual:Pooled) |  | 3.6 | 7.3 |

## 3 Results

### 3.1 Quality control and mapping

All FASTQ files were of high quality with an average per base Phred score >32. Filtering by conversion rate efficiencies resulted in the removal of four *C. fluminea* samples with less than 98.5% conversion efficiency and a further two *D. polymorpha* due to possible over-conversion and poor recovery of the control sequences. All four pooled libraries had adequate conversion rates. For the remaining samples the conversion efficiencies in the CpG context were 99.34 *±* 0.20% for the positive control while maintaining methylation levels of 97 *±* 0.6% on the negative control (Table S3). For the individual libraries, the average number of reads after filtering and end-trimming was 75 *±* 9.3 million ( *μ±* S.D.) for *C. fluminea* and 70 *±* 9.7 million for *D. polymorpha* (Table S1). The pooled libraries had reads slightly under 10x the value of a single individual library; 669 *±* 10 million for *C. fluminea* and 571 *±* 121 million for *D. polymorpha*. The statistical comparison of the read QC measures between pollution treatments within species showed that all groups had a normally distributed number of reads, number of duplicated reads and proportion of aligned reads (p-value > 0.05, Shapiro Wilk Test).

### 3.2 Larger data loss in individual libraries compared with pooled libraries

Here we used the conventional, conservative minimum coverage filter of 10x for all work. Consequently, the minimum resolution of methylation difference that could be detectable is expected to be 10%. While the sequencing effort per individual was the same between individual and pooled libraries, the individual libraries yielded several orders of magnitude fewer sites than the pooled libraries for both species (Table 2). Specifically, 95 - 97% of the individual library data was filtered out at this step, while 30 - 35% of the pooled library data was filtered out. This resulted in approximately 700,000 to 1,200,000 sites per individual library, while we obtained approximately 15 to 32 million sites per pooled library (Table 2). We also observed a far larger proportion of the sites that had low coverage for individual libraries. Sites with high coverage (> 32x) were not well represented in the individual sequenced data (< 10K sites) but are very well represented in the pooled libraries (> 100K sites). Coverage values are variable among individual libraries, but there were no large deviations from the mean to warrant exclusion in all but one treatment group (Table S1).

The difference in analyzed data between individual and pooled libraries is even more striking when looking at the set of 1 kb regions in common among individual libraries (i.e. after the analytical ‘union’ step), that is subsequently used for differential methylation analysis (Table S2). There, we found that only 5,158 and 18,800 regions were common to 75% of the individuals for *Dreissena* and *Corbicula*, respectively. The number of regions in common drops to 1,150 and 7,832 if regions have to be present in each individual. In contrast, the number of regions available for differential methylation analysis was 798,744 and 408,906 for the pooled library data of *Dreissena* and *Corbicula*, respectively. Similarly, the number of regions retrieved from the computationally pooled datasets was 191,075 and 94,234 for *Dreissena* and *Corbicula*, respectively. To summarize, we observed greater data loss in individual vs. pooled library data due to two major steps: 1) initial minimum coverage filtering per individual, and 2) union step to find the sites or regions in common among individuals.

### 3.3 Evaluation of agreement between pooled and individual libraries

#### 3.3.1 Global DNA methylation and correlations of genome-wide methylation levels

*Corbicula fluminea* had slightly lower global methylation levels compared to *Dreissena polymorpha* (16% ± 0.7% vs 19.9% ± 0.6%). Differences in methylation were negligible between the polluted and non-polluted treatments, and between the pooled and individual libraries for both species (*C. fluminea:* individual libraries 16.04% ± 0.77% vs 16.12% ± 0.61%, pooled libraries 16.1% vs 16.2%, *D. polymorpha:* individual libraries 20.01% ± 2.29% vs 19.99% ± 0.63%, pooled libraries 19.5% vs 20%).

We examined the correlation of methylation percentage ( *β*) values between individual libraries and pooled to test for congruence between the two datasets that are expected to be equivalent. Correlations were slightly stronger for *C. fluminea* (0.93 - 1.00) compared to *D. polymorpha* (0.91 - 1.00; Figure 2), but overall similar trends were detected. The *β* values were strongly correlated between individual and pooled libraries with the pairwise correlation coefficients not going below 0.95, irrespective of the pollution treatment, species or loci subset. This is expected if pooled libraries represent an average of the individual libraries and where data have not been centered (see Xu et al. (2015) for the importance of centering).

**Figure 2:**
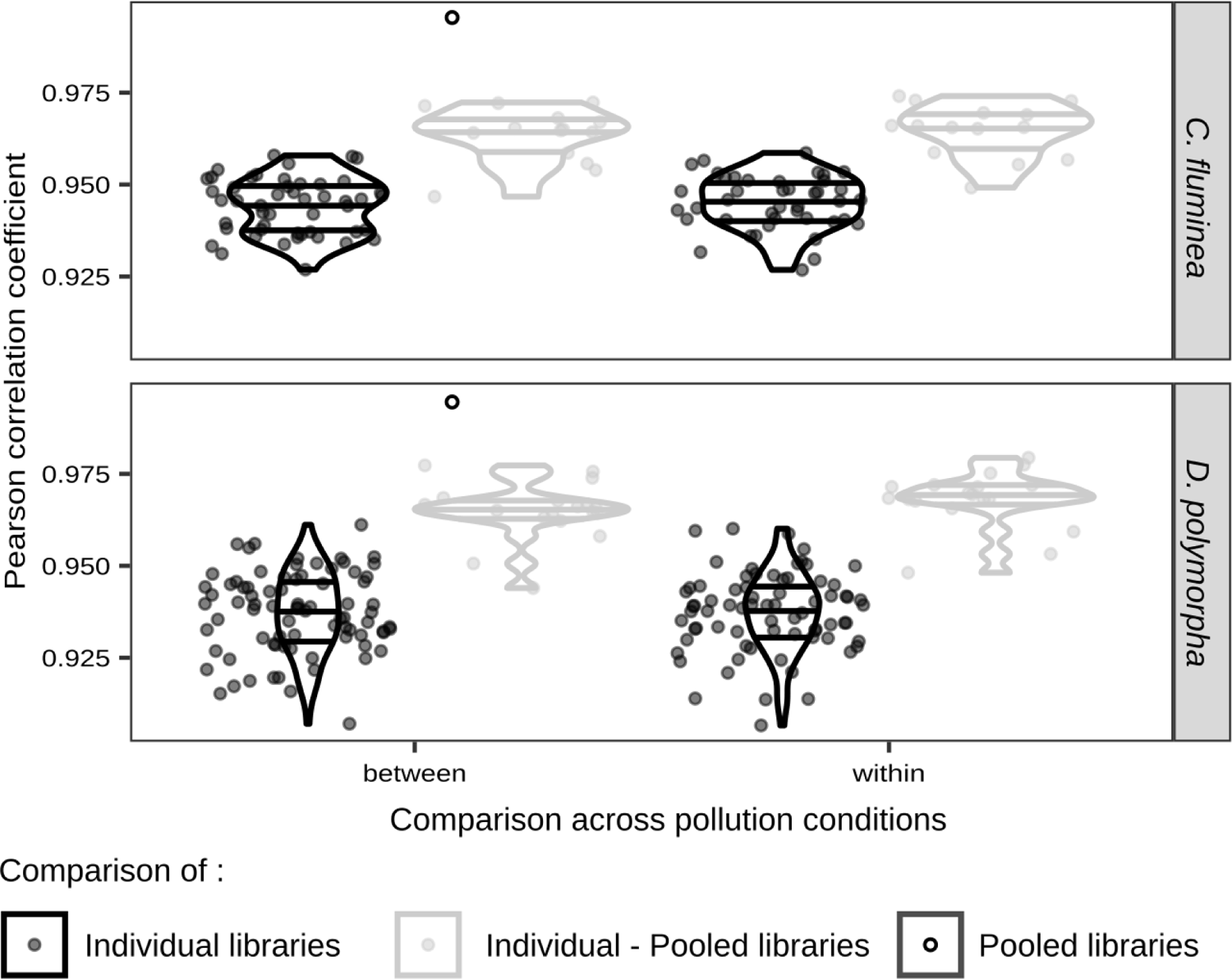
Scatter plots of the correlation coefficients for correlations of percent methylation between pollution treatments. Pearson correlation coefficients are based on the *β* values (per-region % methylation) for each pair of libraries when using all samples.

Unexpectedly, however, individual libraries did not correlate better with pooled libraries of the same treatment group compared to correlations across treatment groups (Figure 2). For both species, pooled libraries correlated best with each other (∼1.00). These results suggest that pooling determines correlation more than pollution treatment, and that there is not a lot of variation in DNA methylation between pollution treatments. For both species, the individual libraries from the pollution treatment had the lowest correlation coefficients for within treatment correlations (Figure 2). This suggests that there may be a pollution-related response in methylation estimates. Overall, we found that genome-wide methylation levels of individual and pooled libraries were well correlated following our expectations.

#### 3.3.2 Agreement among PCA jackknife iterations

To understand the similarity among samples, we tested for clustering using a principal component analysis on the *β* values. We measured heterogeneity of the signal across the genome with a standard error based on a delete-one jackknife. This measured changes to the PCA coordinates when removing a linkage group with each iteration of the PCA. A Tucker’s coefficient was used to gauge similarity across the PCA iterations. For both species and nearly all PCA iterations, the Tucker’s coefficient was greater than 0.95 indicating an overall agreement in signal between the global PCA and each iteration (see Figures S2 & Figures S3). For *C. fluminea*, one of the jackknife iterations gave notably lower Tucker’s coefficients and greater Procrustes D values than the remaining iterations, indicating a disproportionate influence from the respective linkage group. As we only examined the first eight linkage groups in this work, we cannot say if the excluded linkage groups may also be disproportionately influential in the PCAs. The result does argue that some linkage groups may have notable divergences from the majority of the genome. The iteration excluding LG02 produced coefficients below ∼0.95 for the PCAs including data from the polluted treatment (Figures S2). This suggests that LG02 has a significant influence on all other iterations of the PCA and this may be driven more by the polluted treatment group.

#### 3.3.3 PCA of genome-wide methylation levels

In the *C. fluminea* PCA with all samples (94,912 regions; Figure 3A), the proportion of variance captured by each eigenvector declined gradually with the primary vector capturing ∼ 10 % of the variance and the first 11 accounting for top 90 % of the variance. There was a subtle difference between pollution treatments (Figure 3B). There were no clear differences in the variation within either treatment group. The two pooled libraries plotted central to all the individual libraries, supporting that the pooled libraries represent an average of the individual libraries.

**Figure 3:**
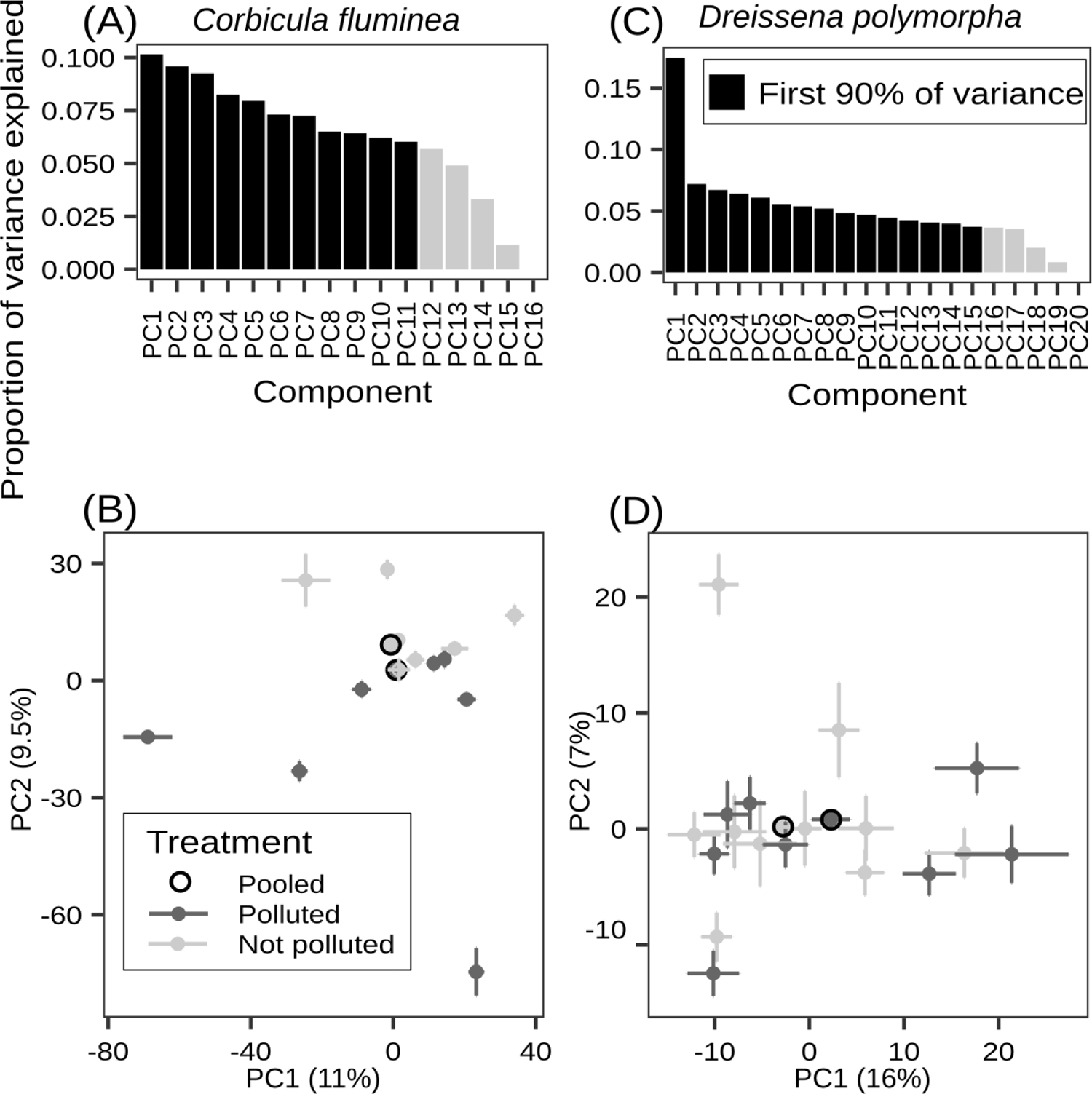
Principal component analysis of the genome-wide percent methylation. Panels A and C show the variance explained by each component for *C. fluminea* and *D. polymorpha*, respectively. Highlighted bars show the components that make up the top 90% of the variance. Panel B and D show the first two components with standard error bars based on the delete-one jackknife in *C. fluminea* and *D. polymorpha*, respectively. The percent variation explained by each axis is indicated on the axis label. The pooled libraries plot centrally to the individual libraries from the same treatment, highlighting that each pooled library represents the average signal of a particular treatment or population.

In the *D. polymorpha* PCA with all samples (190,879 regions; Figure 3C), the first eigenvector captured a significant part of the variance (16%) within the pollution treatment. The remaining vectors captured similar proportions of the variance (∼ 4 - 7 %). There was no clear support for a directional or consistent difference in central tendency of either pollution group (Figure 3D). The two pooled libraries plot central to all the individually-sequenced data, corroborating that the pooled libraries represent an average of the signal found in individual libraries.

#### 3.3.4 Comparison of differential methylation in individual vs. pooled libraries

We used the pollution treatment to assess if differential methylation estimates were similar between individual and pooled libraries. For both species, there were orders of magnitude more DMR in either wet-lab or computationally pooled libraries compared with individual libraries (Figure 4). This is not surprising given the large difference in input data between individual and pooled libraries (see section 3.2). The pooled library of *C. fluminea* produced 943 and 920 DMR (hyper- and hypo-methylated, respectively) while the computationally pooled library produced 707 and 462 DMR (hyper- and hypo-methylated, respectively). In contrast individual libraries produced only one DMR (one per hyper- and hypo-methylated), making the comparison with the pooled dataset not possible.

**Figure 4:**
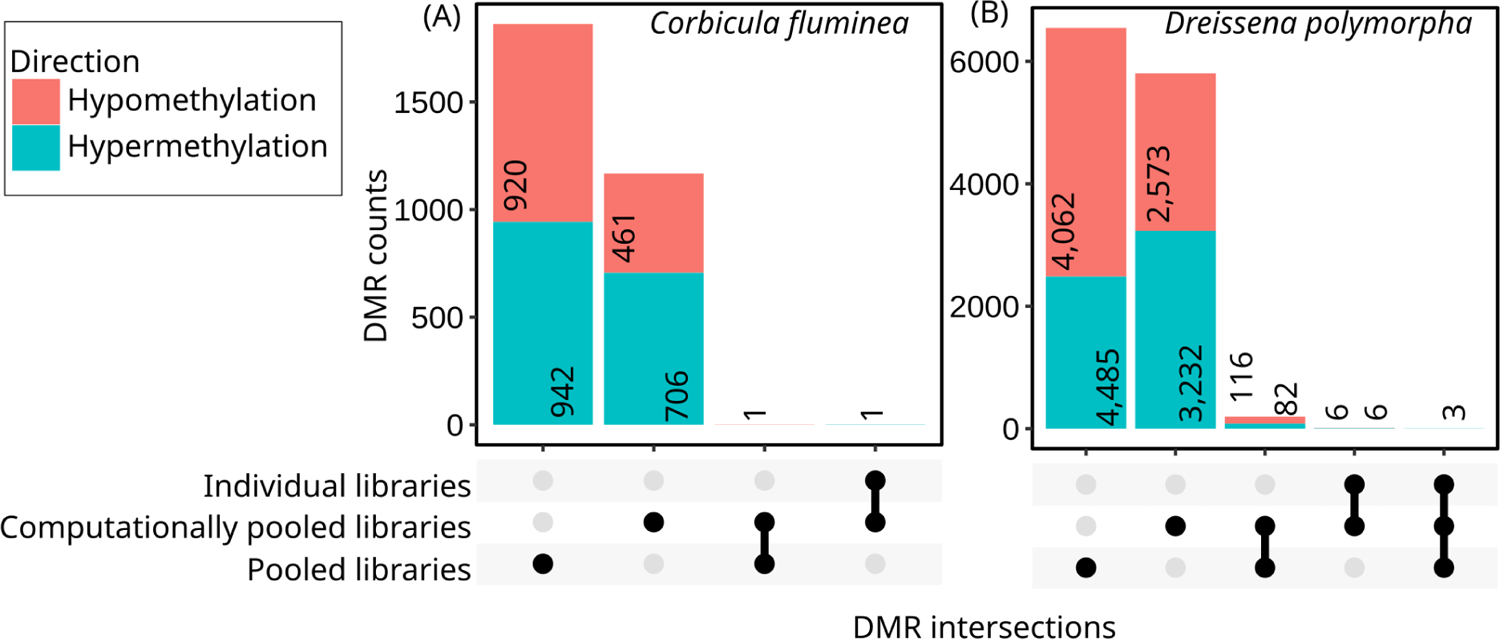
Intersection of the identified DMRs between the polluted and non-polluted localities from the individual, pooled and computationally pooled libraries. The individual, pooled and computationally pooled libraries are each a ‘set’ of DMRs as show by the rows at the bottom. The ‘intersections’ (columns) are the DMRs shared between sets. The dot-plot in the bottom shows how DMRs from each set are distributed among intersections. The central barplot shows the number of DMRs within a particular intersection. Annotations indicate the number of DMRs which are either hyper- or hypo-methylated.

Similarly, the pooled library datasets of *D. polymorpha* produced 2,570 and 4,178 (hyper- and hypo-methylated, respectively), the computationally pooled library produced 3,323 and 2,695 (hyper- and hypo-methylated, respectively). Individual libraries produced 9 and 6 (hyper- and hypo-methylated, respectively) (Figure 4). It is worth noting that each DMR from individual library datasets was also in common with the DMRs of either the pooled library or computationally pooled library dataset.

Despite the large number of regions detected by the tests with the pooled and computationally pooled data, there was low overlap in the identified regions. Only one DMR was identified by the pooled and computationally pooled datasets in *C. fluminea* and only 201 DMR in *D. polymorpha*. In both species this amounted to < 3.5% of the regions from either test. This supports a predominant incongruence in the results between the pooled and computationally pooled datasets. Furthermore, the relative proportion of hyper- to hypo-methylated DMR changed between tests. For *C. fluminea,* computationally pooled data produced a greater hyper-methylated region: hypo-methylated region ratio (0.23 : 0.15) compared with the pooled data (0.31 : 0.30). For *D. polymorpha*, both pooled data produced unequal proportions, but a greater hyper-methylated region: hypo-methylated region ratio was observed for computationally pooled libraries (0.26 : 0.20) compared to the pooled data (0.20 : 0.32).

### 3.4 Estimates of the required sampling effort and the recovered power

To assess the available power in our data, we estimated the level of power recovered with our current sampling effort and the sampling effort needed to achieve power at a level of 80% at each region. We used locus-specific estimates of effect size and mean difference in methylation (MDM) between treatment groups (i.e. polluted vs. non-polluted).

A low proportion of regions were deemed invariant. The pooled and computationally pooled libraries produced more invariant regions than the individual libraries (Table S4). For the variant regions (i.e. MDM > 0), the sampling effort estimates were very large. For the individual libraries, estimates were predominantly more than 100 per treatment group (Figure 5A & D) and we found that no loci achieved sufficient power with our current sampling effort. Regions with sample effort estimates below 100 were only ∼7.5% (98) - 14.7% (1,506) of the total loci.

**Figure 5:**
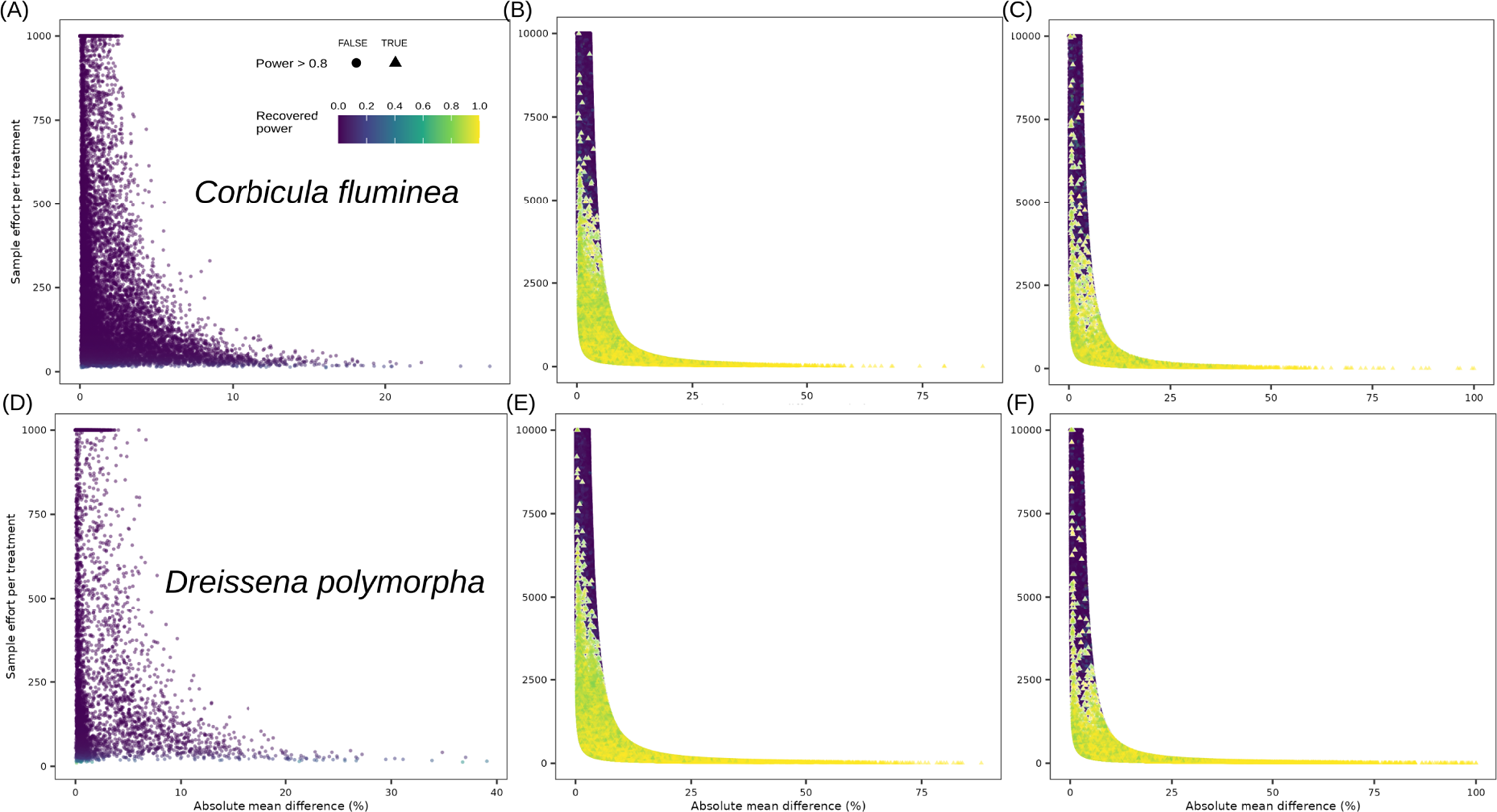
Estimates of the required per-locus sampling effort and the achieved power. Panel A - C shows the estimates for *C. fluminea* and panel D - F shows the estimates for *D. polymorpha* with separate plots for individual (panels A D), pooled (panels B E) and computationally pooled libraries (panels C F). Dots are individual loci and colours indicate the estimated power achieved. Note that in panels A D loci with sampling effort estimates > 1000 were capped to 1000 and in panels B, C, E F, loci with sampling effort estimates > 10,000 were capped to 10,000. Sampling effort is measured as the number of biological replicates per treatment for individual libraries in panel A D and sequencing coverage per pooled treatment for pooled libraries in panel B, C, E F. Loci with power > 0.8 were plotted above other loci to aid visibility.

Similarly, in the pooled and computationally pooled data, estimates of required sampling effort were >1,000x and almost exclusively >100x for both species (Figure 5 B, C, E, F). For a set of 10 individuals, this would require more than 100x coverage each to achieve adequate coverage for less than 50% of the variance distribution. We recovered adequate power at only ∼ 7 - 16 % of regions in the pooled and computationally pooled libraries, highlighting that the majority of our regions were under-powered. Computationally pooled libraries produced a greater proportions of regions with coverage estimates below 100x compared to pooled libraries; 3 % (2,432) vs 0.5% (2,901) for *C. fluminea* and 8 % (12,128) vs 1% (8,999) regions for *D. polymorpha*, (Figure 5 B & E).

Across all comparisons, regions with the lowest MDM had the lowest power and largest sampling effort, which is to be expected as larger differences require fewer samples. Nearly the entire MDM distribution was predominantly under-powered. For example, the lowest 50% of the MDM distribution resulted in the following power values: MDM < 0.52 % had power < 0.1 for individual libraries (Figure 5 A & D), MDM< 1.5 % had power < 0.2 for pooled *C. fluminea* data, pooled *D. polymorpha* (Figure 5 B & E), and computationally pooled data from *C. fluminea* (5 E) and MDM < 2.4 % had power < 0.2 for *D. polymorpha* computationally pooled data (Figure 5 F). The upper 50 % of MDM distribution had median power < 0.8 (Table S5). Only the highest 10% of the MDM distribution for the computationally pooled *D. polymorpha* data had a median power > 0.8; ∼ 16.3 % of the total regions. In contrast, pooled libraries produced sufficient power at only 7 % (*C. fluminea*) and 15 % (*D. polymorpha*) of regions. Computationally pooled *C. fluminea* data produced only 8.6 % regions with sufficient power. These loci covered an MDM range of 4 - 40% for individual libraries, 7 - 88% for pooled libraries and 11 - 100% for computationally pooled libraries.

With regards to differences between the species, we see that *D. polymorpha* had a greater proportion of regions with achievable coverage, this being double the proportions for *C. fluminea* in the pooled data and computationally pooled data. For the individual libraries this proportion was only half *C. fluminea*. Similarly, a greater proportion of regions achieved sufficient power for *D. polymorpha*.

Finally we also note the greater MDM for *D. polymorpha,* as much as 39 % for individual libraries (vs 18 % *C. fluminea*). For the pooled and computationally pooled data, the two species had equal ranges for MDM, 88% and 100%. These differences may be attributed to sample sizes between the species and the possible differences in natural variation.

## 4. Discussion

### 4.1 Individual and pooled libraries provide similar genome-wide methylation estimates

Here we examined if cost-effective pooled whole-epigenome libraries provide equivalent biological results to individually sequenced libraries. We found that pooled libraries produced a congruent epigenetic signature with individual libraries at the genome-wide level as seen with pairwise correlations and a PCA. Our estimates of global methylation were also remarkably stable between pooled and individual libraries. These results are in line with previous research supporting a global or genome-wide correspondence of DNA methylation levels from pooled and individual libraries (Docherty et al., 2009; Docherty et al., 2010; Gallego-Fabrega et al., 2015). In our data both pooled and individual libraries showed a negligible difference in methylation between the polluted and non-polluted treatment groups and for both species but in all cases DMRs were detected.

### 4.2 Pooled libraries provide more data than individual libraries while reducing costs

We found that there is at least a 7-fold decrease in the cost per sample for gross pooled libraries compared with individual libraries (Table 3). The reduction depends on the number of biological replicates in each pool and this allows a great degree of flexibility to balance sample sizes, coverage and cost. The pooled libraries also produced orders of magnitude more final regions (between 60 – 600x more than individual libraries) and lost notably less data throughout the workflow. When estimating the required sampling efforts to reach sufficient power per locus, pooled libraries produced as much as 30x more regions with an achievable estimate (98 vs 3,000 regions for *C. fluminea*). Achievable is defined here as regions with estimates of <100 samples per treatment group (individual libraries) or coverage estimates below 100x (pooled libraries). The achieved power was greater for the pooled libraries, as much as 16% of regions had sufficient power (power = 0.8), while the individual libraries for both species had no regions reach the threshold. All of this strongly supports that pooled libraries produce sufficient data more reliably compared with individual libraries given equal per-sample coverage. In practice, our pooled libraries detected >1,000 DMRs while the individual libraries detected <15 DMRs. These differences will make a meaningful impact on the return of investment and the possible scope of downstream interpretation in research. We also employed computational pooling which pools the data of the individual libraries *post-hoc*. Here we found it to successfully mitigate much of the data loss that individual libraries suffered from during the QC process. Pooled and computationally pooled libraries produced comparable volumes of input data and DMRs detected.

### 4.3 Pooled and computationally pooled libraries provided different DMR in our dataset

Beyond the global signal, our results showed that genome-wide congruence does not necessarily imply corresponding DMR signals for pooled and individual libraries. We found low overlap in DMR which is contrasting with the correlations and PCA. While the difference in volume of input data played a significant role for individually sequenced libraries, the results from the computationally pooled libraries also showed that the discrepancy is not solely due to the number of regions detected.

In our dataset, pooled and computationally pooled libraries both had several thousand DMRs detected yet had < 4% regions in common. These results argue that there are discrepancies in the underlying signal between the wet-lab pooled and computationally pooled libraries. These differences may arise from several factors: 1) the individuals compared between the pooled and computationally pooled libraries were not exactly the same as four *Corbicula* and two *Dreissena* individuals were excluded from downstream analyses as they failed conversion rate quality control; 2) lack of normalization of individual data before computational pooling; 3) stochasticity in the library preparation and sequencing processes (e.g. differential PCR during library preparation; cryptic biases in sequencing among the specimens of the pooled libraries). Based on these results, we discuss below the benefits and drawbacks of sample pooling, as well as possible improvements and ways forward.

### 4.4 Benefits and drawbacks of sample pooling and recommendations

WepiGS offers important opportunities for ecological and evolutionary studies, hence it is crucial to optimally use resources and consider trade-offs before initiating a project. We emphasize that there is not a single optimal solution for all projects and that the decision to sequence pooled or individual libraries depends on the scientific question of a particular project and should be planned at very early stages. Here we put our work in context and provide a summary of the key benefits and drawbacks of pooling libraries for epigenome sequencing (WepiGS), as well as their implications (summary in Table 4).

**Table 4:** Benefits and drawbacks of DNA pooling before library preparation.

| Topics | Implications | Libraries |  |
| --- | --- | --- | --- |
|  |  | Individual | Pooled |
| Costs | Higher wet lab costs for individual libraries. Cost savings can be adjusted according to the pooling scheme (See Table 3 for details) | - | + |
| Power | Increased number of individuals included in a pool improves accuracy of population-level metrics (e.g. Response to treatments, differences between environmental conditions) and increases power to detect differences | - | + |
| Power / costs | 20-30-fold more data when pooled libraries are sequenced at an equivalent sequencing effort to the individual libraries (see Table 2 for details). Additional cost savings are possible if sequencing effort of pooled libraries is reduced. | - | + |
| Flexibility | Individual information (covariates) cannot be used with pooling. Nested pooling (pooling by condition, e.g. sampling locality, sex, age, tissue, experimental condition, ...) is needed to measure variability in the data. Data reuse for subsequent projects is challenging | + | - |
| Data analyses | Differences in individual conversion rates or individual sequencing depth are not taken into account when samples are pooled. Possible biased representation of some samples in the pool cannot be accounted for. | + | - |
| Data analyses | Greater computational resources needed resulting from greater data volumes of pooled libraries. Many tools for methylation analyses are not adapted to handle large datasets from pooled libraries. | + | - |
| Data analyses | Fewer, less flexible statistical tests are available for pooled datasets. | + | - |

Starting with the benefits, we showed that pooled libraries can be up to 7-fold more cost effective than individual libraries, when comparing wet lab costs. These costs are likely to be a limiting factor into the future. There is flexibility in cost adjustment when the pooling scheme varies (e.g. deciding how many pools to prepare) but there are presently limited research on the trade-offs of different degrees of pooling. Another important benefit of pooled libraries is that the number of individuals per pool can be increased; typically the number of individuals per investigated population is between 10 and 20, however it has been shown that a larger number of individuals is required to achieve sufficient power in natural populations (Tsai & Bell, 2015, Lea et al 2017). For instance, when a predictor variable explains 15% of the difference between populations, 125 individuals per population are needed to reach 50% power (Lea et al, 2017). A third advantage that we did not expect was that a larger proportion of the sequencing data can be used with pooled libraries, resulting in a 20-30 fold increase in retrieved loci in our dataset. This was not only due to the higher coverage of pooled libraries, but also because there was a single union step using the pooled datasets (i.e. finding the loci in common among libraries, which typically leads to a large loss of data). Together with this, if a high sequencing coverage is not necessary (e.g. in our case 100x per pooled library), researchers can decide to lower the sequencing effort per pooled library, possibly decreasing even more the project costs.

Using pooled libraries has several drawbacks, though, the most important one being that there is no possibility of going back to the individual data. Hence, researchers should be extremely careful when thinking about the pooling design, and make sure that every covariate that may impact the signal in the data has been taken into account (e.g. sampling locality, sampling time in the year, sex, age, tissue, experimental condition, etc.). If these covariate can be clearly identified and separated in sub-pools, then pooling the DNA of samples may be a good option to increase power and decrease costs. In contrast, if covariates cannot be identified or if the variability in the data is not known (e.g. first epigenomic experiment), we would recommend against pooling.

In addition, individual libraries provide more flexibility and higher resolution as groups and comparisons can be done *a posteriori* (e.g. testing the impact of different covariates in a pilot study) and data can be reused for future projects (e.g. adding individuals from different populations or time points, or different comparisons can be made). Other drawbacks of using pooled libraries arise from the data analysis side. For instance, individual samples that failed cannot be excluded (e.g. low conversion rate; low amount of sequencing data), and thus equal conversion rates and sequencing depth for all individuals in a pool is assumed. Furthermore, we observed that a large amount of computational resources was required to process the pooled datasets (e.g. alignment, methylation calling) and that the currently widely used bioinformatic tools are not well adapted to handle such large datasets (e.g. MethylKit running in R). Researchers could decide to filter out invariant sites early in the data analysis, and/or decrease the sequencing depth of the pools (i.e. less than 10x per sample), however the minimum sequencing depth to obtain meaningful population methylation rates is not known. We note that this issue arises when study organisms have large genomes (in our case 1.6 Gb and 1.8 Gb), but it may be less of a problem for organisms with smaller genome sizes (e.g. less than 1 Gb). Finally, we also noticed that there are fewer and less flexible statistical tests available for data analysis (e.g. Logistic regression cannot be used with two samples). We hope that new tools that can handle large pooled epigenomic datasets will be developed in the future.

To conclude, because individual libraries provide greater flexibility and control, they are the best option to explore a first epigenomic dataset where covariate variation is unknown, or when samples are rare or limited. However, we believe there are situations in which pooling DNA before library preparation would be the best option to obtain population-level signals (Futschik & Schlotterer, 2010, Kaplow et al., 2015), increase power and decrease costs. For instance, when the number of individuals per population is not limited (e.g. abundant species), in well-studied systems where epigenomic variation is already known and researchers want to increase power in follow-up studies. In these systems, either a clear separation of covariates is possible, or organisms are small and whole organisms are used for DNA extraction, making sure that all covariates are captured in a single DNA extraction (Harney et al., 2022). Finally, pooling would be particularly well-suited in systems with small genome sizes to facilitate downstream analyses.

### 4.5 Possible improvements and ways forward

We have shown that pooled libraries provide estimates of genome-wide methylation comparable to individual libraries. However, signals of differential methylation at specific regions were not congruent between individual and pooled libraries, mostly as a result of large differences in the number of loci retrieved and the power. Even when pooling computationally the data of individual libraries, the overlap with the empirically pooled datasets was low. This may arise from the fact that six individuals were excluded from the analyses due to low conversion rates and low sequencing data, resulting in actual differences between the pooled and computationally pooled datasets. In addition, stochasticity in the library preparation and sequencing processes may have led to further discrepancies between these datasets. Together with this, we worked on two empirical systems with relatively low global DNA methylation levels (16-20%), which likely explains the low power recovered in our datasets. It is possible that when working with species with higher global DNA methylation levels there would have been more congruence between DMR of pooled and computationally pooled datasets.

In addition, these epigenomic datasets were the first ones for the two species of interest, *Corbicula fluminea* and *Dreissena polymorpha*. Therefore, global DNA methylation levels were previously unknown, as well as the level of covariate variation. Based on these results, before considering pooling, we would recommend to perform a pilot study using individual libraries to assess these metrics and make an informed decision about pooling individuals in subsequent studies or not. Furthermore, we acknowledge that we did not perform simulations in this study, because we wanted to focus on empirical data to explore commonalities and differences between individual and pooled libraries produced in the lab. Thus, we aimed to obtain a very practical result close to a real experiment. In future studies, it would be interesting to simulate the minimum coverage of a pool required to obtain reliable population-level DNA methylation rates, as a way of facilitating downstream analyses and further decreasing project costs. To conclude, our study brings important insights on the relevance of pooling DNA of individuals before library preparation in epigenomic studies of natural populations, and we believe that it will help researchers in making informed decisions for future epigenomic projects.

## Acknowledgments

The project was funded by the Swiss Federal Institute of Aquatic Science and Technology (Eawag). We also thank Niklaus Zemp for bioinformatic support. Sequencing was performed at the Functional Genomics Center Zürich (FGCZ), Switzerland. Data analyses were performed at the Euler High - Performance Computer Centre in Zürich, Switzerland (https://sis.id.ethz.ch/services/hpc/). Data produced and analyzed in this paper were generated in collaboration with the Genetic Diversity Centre (GDC), ETH Zürich. RJD was supported by the National Research Foundation of South Africa. SS received funds from Short Term Scientific Missions provided by the COST Action “Conservation of freshwater mussels: a pan-European approach” (CONFREMU) CA1823. The authors declare no conflict of interests.

## Data Accessibility and Benefit - Sharing

**Genetic data:** Raw sequence reads are deposited in the Sequence Read Archive (https://www.ncbi.nlm.nih.gov/sra) (To be deposited) **Metadata:** Related metadata can be found in the EAWAG internal database, ERIC (To be deposited). Benefits Generated: Benefits accrue provides unique sample identifier tags that can be matched to both the deposited genetic data and deposited metadata from the sharing of data and results publicly.

## Author Contributions

RJD – Analyses, Writing; BSM – Supervision, Writing; SS, MG – Laboratory work, Writing CDT-Design, Sample contribution, NR - Design, Sample contribution, SS – Design, Sample contribution; AATW – Design, Supervision, Writing, Funding

## Supplemental Tables

**Table S1:** Summary of the paired sequence data used for alignment. Presented for each library are the estimates of the total number of paired reads, the proportion of reads successfully aligned, the estimated levels of duplication, the resultant number of Cytosines in the CpG context and the coverage at CpG sites. Samples removed from downstream analyses due to low conversion efficiencies are indicated in grey. The average across treatment and pooling scheme are indicated at the bottom of each set of samples. The averages do not include samples removed from downstream analyses. S.D. = Standard deviation.

| Species | Sample | Paired reads |  |  |  |  |  | Cytosine |  |  |  |
| --- | --- | --- | --- | --- | --- | --- | --- | --- | --- | --- | --- |
|  |  | Total | S.D. | % aligned | S.D. | % duplication | S.D. | Total | S.D. | Coverage | S.D. |
| *Corbicula fluminea* | C10P | 79,904,532.00 |  | 34.18 |  | 20.48 |  | 973,770,066.00 |  | 4.14 |  |
|  | C1P | 74,738,403.00 |  | 32.03 |  | 20.5 |  | 887,207,192.00 |  | 3.79 |  |
|  | C2P | 84,759,893.00 |  | 34.09 |  | 18.57 |  | 1,051,349,940.00 |  | 4.37 |  |
|  | C3P | 65,165,994.00 |  | 34.37 |  | 19.01 |  | 778,022,325.00 |  | 3.26 |  |
|  | C4P | 38,398,067.00 |  | 34.35 |  | 16.05 |  | 487,725,519.00 |  | 1.96 |  |
|  | C5P | 68,910,145.00 |  | 34.73 |  | 16.84 |  | 825,628,926.00 |  | 3.36 |  |
|  | C6P | 75,235,776.00 |  | 33.54 |  | 18.3 |  | 922,937,770.00 |  | 3.79 |  |
|  | C8P | 55,808,379.00 |  | 34.76 |  | 17.24 |  | 682,092,108.00 |  | 2.77 |  |
|  | C9P | 83,025,026.00 |  | 34.05 |  | 16.99 |  | 1,059,124,248.00 |  | 4.29 |  |
|  | Average | 71,826,893.57 | 9,311,387.23 | 33.95 | 0.94 | 18.48 | 1.57 | 875,540,376.43 | 125,935,097.78 | 3.63 | 0.53 |
|  | C10NP | 79,616,730.00 |  | 34.33 |  | 19.63 |  | 964,245,233.00 |  | 4.06 |  |
|  | C1NP | 92,972,078.00 |  | 34.31 |  | 19.15 |  | 1,152,249,279.00 |  | 4.78 |  |
|  | C2NP | 83,972,229.00 |  | 33.25 |  | 21.27 |  | 992,946,861.00 |  | 4.31 |  |
|  | C3NP | 75,932,743.00 |  | 33.9 |  | 17.96 |  | 942,277,154.00 |  | 3.86 |  |
|  | C4NP | 64,381,899.00 |  | 34.67 |  | 18.09 |  | 809,494,157.00 |  | 3.33 |  |
|  | C5NP | 78,291,875.00 |  | 34.74 |  | 18.99 |  | 984,091,085.00 |  | 4.1 |  |
|  | C6NP | 74,741,855.00 |  | 33.71 |  | 18.07 |  | 917,716,701.00 |  | 3.76 |  |
|  | C8NP | 74,770,619.00 |  | 34.52 |  | 17.87 |  | 926,653,920.00 |  | 3.81 |  |
|  | C9NP | 82,647,061.00 |  | 33.73 |  | 20.27 |  | 975,358,391.00 |  | 4.14 |  |
|  | Average | 78,055,450.43 | 8,880,630.31 | 34.1 | 0.5 | 18.86 | 1.26 | 957,940,472.14 | 103,198,480.14 | 3.99 | 0.46 |
|  | CpoolP | 661,904,290.00 |  | 34.53 |  | 29.46 |  | 7,004,096,743.00 |  | 33.62 |  |
|  | CpoolNP | 675,878,620.00 |  | 34.08 |  | 31.4 |  | 7,001,527,240.00 |  | 34.51 |  |
|  | Average | 668,891,455.00 | 9,881,343.51 | 34.31 | 0.31 | 30.43 | 1.37 | 7,002,811,991.50 | 1,816,913.00 | 34.06 | 0.63 |
| *Dreissena polymorpha* | D10P | 65,410,675.00 |  | 49.59 |  | 15.48 |  | 1,160,699,600.00 |  | 4.09 |  |
|  | D1P | 73,460,038.00 |  | 49.4 |  | 15.07 |  | 1,349,558,024.00 |  | 4.74 |  |
|  | D2P | 59,121,375.00 |  | 48.13 |  | 14.39 |  | 1,039,914,761.00 |  | 3.63 |  |
|  | D3P | 68,703,649.00 |  | 48.54 |  | 16.13 |  | 1,269,970,624.00 |  | 4.51 |  |
|  | D4P | 51,119,921.00 |  | 47.23 |  | 14.32 |  | 872,596,859.00 |  | 3.02 |  |
|  | D5P | 64,776,750.00 |  | 51.77 |  | 13.92 |  | 1,263,320,184.00 |  | 4.34 |  |
|  | D6P | 42,903,550.00 |  | 49.12 |  | 15.47 |  | 751,604,109.00 |  | 2.64 |  |
|  | D7P | 47,844,529.00 |  | 48.37 |  | 14.33 |  | 841,215,204.00 |  | 2.94 |  |
|  | D8P | 66,746,309.00 |  | 48.32 |  | 13.67 |  | 1,227,896,716.00 |  | 4.21 |  |
|  | D9P | 63,351,525.00 |  | 50.2 |  | 14.39 |  | 1,203,155,822.00 |  | 4.18 |  |
|  | Average | 64,086,280.25 | 6,674,156.69 | 49.15 | 1.42 | 14.67 | 0.83 | 1,173,389,073.75 | 151,544,719.09 | 4.09 | 0.54 |
|  | D10NP | 75,884,193.00 |  | 47.74 |  | 15.73 |  | 1,278,999,481.00 |  | 4.51 |  |
|  | D1NP | 79,359,990.00 |  | 48.92 |  | 14.97 |  | 1,449,986,916.00 |  | 5.08 |  |
|  | D2NP | 69,143,806.00 |  | 49.43 |  | 14.77 |  | 1,289,478,957.00 |  | 4.51 |  |
|  | D3NP | 72,289,318.00 |  | 49.33 |  | 15.08 |  | 1,354,203,964.00 |  | 4.77 |  |
|  | D4NP | 72,492,019.00 |  | 47.68 |  | 15.09 |  | 1,338,539,670.00 |  | 4.68 |  |
|  | D5NP | 73,438,833.00 |  | 49.23 |  | 16.04 |  | 1,343,852,759.00 |  | 4.79 |  |
|  | D6NP | 71,852,551.00 |  | 49.77 |  | 13.93 |  | 1,351,093,249.00 |  | 4.69 |  |
|  | D7NP | 73,660,941.00 |  | 48.89 |  | 15.49 |  | 1,342,094,303.00 |  | 4.75 |  |
|  | D8NP | 98,739,072.00 |  | 48.89 |  | 15.95 |  | 1,751,923,420.00 |  | 6.23 |  |
|  | D9NP | 66,962,735.00 |  | 49.6 |  | 13.95 |  | 1,258,500,785.00 |  | 4.38 |  |
|  | Average | 75,382,345.80 | 8,872,805.68 | 48.95 | 0.72 | 15.1 | 0.74 | 1,375,867,350.40 | 142,273,247.94 | 4.84 | 0.53 |
|  | DpoolP | 657,309,352.00 |  | 48.94 |  | 24.88 |  | 10,418,395,650.00 |  | 41.31 |  |
|  | DpoolNP | 485,624,776.00 |  | 49.2 |  | 19.65 |  | 8,590,178,928.00 |  | 31.9 |  |
|  | Average | 571,467,064.00 | 121,399,327.91 | 49.07 | 0.18 | 22.27 | 3.7 | 9,504,287,289.00 | 1,292,744,441.60 | 36.6 | 6.65 |

**Table S2:** Number of loci retained after filtering for union across libraries. Presented are the mean and standard deviation (S.D.) of the number of regions retained after uniting samples for individual, pooled and computationally pooled libraries. The data loss (% loss) is estimated relative to the mean number of regions before uniting. For individual libraries, we present the results for when using a 75 % and 100 % overlap across libraries.

| Species | Treatment subset | Overlap | Statistical test | Post-union regions | % loss |
| --- | --- | --- | --- | --- | --- |
| *C. fluminea* | Individual libraries | 0.75 | Logistic Regression | 18,800 | 0.79 |
|  |  | 1 | Logistic Regression | 7,832 | 0.96 |
|  | Pooled libraries | 1 | Fisher's Exact Test | 482,906 | 0.08 |
|  | Computationally pooled libraries | - | Fisher's Exact Test | 94,234 | -1.08 |
| *D. polymorpha* | Individual libraries | 0.75 | Logistic Regression | 5,158 | 0.98 |
|  |  | 1 | Logistic Regression | 1,150 | 0.99 |
|  | Pooled libraries | 1 | Fisher's Exact Test | 798,744 | 0.1 |
|  | Computationally pooled libraries | - | Fisher's Exact Test | 191,075 | -1.42 |

**Table S3:** Estimates of the conversion rate efficiencies. Estimates based on the methylation of two spike-in controls. The pUC19 sequence is CpG methylated (negative control) reflecting 99.5 % efficiency when 96 - 98 % methylated. The lambda sequence is entirely unmethylated (positive control) reflecting 99.5 % efficiency when 0.5 % methylated. Parameters with suspect values are highlighted in grey. Samples removed from downstream analyses indicated in **bold**.

| Control |  | pUC19 |  |  |  |  |  |  |  |  |  | Lambda |  |  |  |  |  |  |  |  |  |
| --- | --- | --- | --- | --- | --- | --- | --- | --- | --- | --- | --- | --- | --- | --- | --- | --- | --- | --- | --- | --- | --- |
| Sample Code | Total C | S.D. | C Coverage | S.D. | % CpG | S.D. | % CHG | S.D. | % CHH | S.D. | Total C | S.D. | C Coverage | S.D. | % CpG | S.D. | % CHG | S.D. | % CHH | S.D. | Excluded |
| C1P | 11,350.00 |  | 20.25 |  | 98.1 |  | 1.7 |  | 1 |  | 212,158.00 |  | 22 |  | 0.7 |  | 0.9 |  | 0.9 |  |  |
| **C2P** | 5,762.00 |  | 9.7 |  | 99.5 |  | 21.1 |  | 22.1 |  | 115,737.00 |  | 11.75 |  | 18.8 |  | 24.7 |  | 25.7 |  | x |
| C3P | 6,051.00 |  | 10.11 |  | 97.7 |  | 3.3 |  | 1.7 |  | 97,711.00 |  | 10.11 |  | 1.1 |  | 1.4 |  | 1.5 |  |  |
| **C4P** | 3,187.00 |  | 5.26 |  | 97.5 |  | 7.8 |  | 7.4 |  | 42,339.00 |  | 4.15 |  | 2 |  | 3.5 |  | 3.8 |  | x |
| C5P | 5,236.00 |  | 9.27 |  | 97.8 |  | 2.9 |  | 2.2 |  | 70,070.00 |  | 7.08 |  | 0.7 |  | 1.3 |  | 1.4 |  |  |
| C6P | 11,882.00 |  | 21.54 |  | 97.8 |  | 1.8 |  | 1.5 |  | 213,506.00 |  | 22.32 |  | 0.4 |  | 0.6 |  | 0.6 |  |  |
| C8P | 3,534.00 |  | 5.77 |  | 99.5 |  | 2.3 |  | 1.3 |  | 88,186.00 |  | 8.94 |  | 0.6 |  | 0.7 |  | 0.9 |  |  |
| C9P | 11,382.00 |  | 19.95 |  | 99.1 |  | 2.6 |  | 1.6 |  | 208,825.00 |  | 21.41 |  | 0.7 |  | 0.9 |  | 0.9 |  |  |
| C10P | 9,158.00 |  | 15.41 |  | 98.9 |  | 3.5 |  | 2.2 |  | 160,481.00 |  | 16.46 |  | 1 |  | 1.4 |  | 1.3 |  |  |
| All P | 7,504.67 | 3,471.67 | 13.03 | 6.37 | 98.43 | 0.81 | 5.22 | 6.23 | 4.56 | 6.86 | 134,334.78 | 66,112.62 | 13.8 | 6.93 | 2.89 | 5.98 | 3.93 | 7.84 | 4.11 | 8.15 |  |
| C1NP | 11,625.00 |  | 19.83 |  | 97.6 |  | 2.5 |  | 1.8 |  | 231,651.00 |  | 23.72 |  | 0.6 |  | 0.7 |  | 0.7 |  |  |
| C2NP | 17,938.00 |  | 31.9 |  | 97.1 |  | 4.2 |  | 3.7 |  | 255,942.00 |  | 26.37 |  | 1.3 |  | 2 |  | 2.1 |  |  |
| C3NP | 13,546.00 |  | 24.55 |  | 97.7 |  | 2.5 |  | 1.8 |  | 201,767.00 |  | 20.59 |  | 0.6 |  | 1.1 |  | 1.1 |  |  |
| C4NP | 9,533.00 |  | 16.12 |  | 98.1 |  | 2.4 |  | 2.1 |  | 156,840.00 |  | 16.13 |  | 0.7 |  | 1.2 |  | 1.2 |  |  |
| **C5NP** | 11,872.00 |  | 21.08 |  | 98.8 |  | 18.2 |  | 19.7 |  | 215,116.00 |  | 21.88 |  | 14.3 |  | 18.6 |  | 18.7 |  | x |
| C6NP | 12,641.00 |  | 22.49 |  | 97.7 |  | 2.3 |  | 1.5 |  | 221,986.00 |  | 23.26 |  | 0.6 |  | 0.7 |  | 0.8 |  |  |
| C8NP | 10,990.00 |  | 18.53 |  | 98.3 |  | 2.3 |  | 1.7 |  | 184,530.00 |  | 18.67 |  | 0.5 |  | 0.7 |  | 0.8 |  |  |
| **C9NP** | 7,640.00 |  | 12.8 |  | 99.4 |  | 13.3 |  | 14.1 |  | 146,817.00 |  | 15.03 |  | 10.7 |  | 14.3 |  | 15.3 |  | x |
| C10NP | 10,539.00 |  | 17.27 |  | 98.3 |  | 1.7 |  | 1.3 |  | 174,080.00 |  | 17.84 |  | 0.6 |  | 0.8 |  | 0.7 |  |  |
| All NP | 11,813.78 | 2,875.67 | 20.51 | 5.52 | 98.11 | 0.69 | 5.49 | 5.98 | 5.3 | 6.76 | 198,747.67 | 36,115.37 | 20.39 | 3.77 | 3.32 | 5.29 | 4.46 | 6.9 | 4.6 | 7.09 |  |
| CpoolP | 48,915.00 |  | 91.82 |  | 97.5 |  | 2.2 |  | 1.6 |  | 937,629.00 |  | 106.48 |  | 0.7 |  | 0.9 |  | 0.9 |  |  |
| CpoolNP | 76,964.00 |  | 157.63 |  | 98 |  | 3.1 |  | 2.4 |  | 1,361,544.00 |  | 161.82 |  | 0.9 |  | 1.3 |  | 1.3 |  |  |
| All pooled | 62,939.50 | 19,833.64 | 124.72 | 46.54 | 97.75 | 0.35 | 2.65 | 0.64 | 2 | 0.57 | 1,149,586.50 | 299,753.17 | 134.15 | 39.13 | 0.8 | 0.14 | 1.1 | 0.28 | 1.1 | 0.28 |  |
| D1P | 13,170.00 |  | 23.47 |  | 98.2 |  | 2.1 |  | 1.2 |  | 218,117.00 |  | 22.29 |  | 0.7 |  | 0.7 |  | 0.7 |  |  |
| D2P | 8,639.00 |  | 15.68 |  | 97.7 |  | 1.9 |  | 1 |  | 137,280.00 |  | 13.75 |  | 0.3 |  | 0.4 |  | 0.5 |  |  |
| D3P | 16,471.00 |  | 29.34 |  | 97.8 |  | 2.2 |  | 1.5 |  | 257,445.00 |  | 26.88 |  | 0.6 |  | 0.8 |  | 0.7 |  |  |
| D4P | 8,228.00 |  | 13.65 |  | 97.9 |  | 1.4 |  | 1 |  | 141,537.00 |  | 14.29 |  | 0.7 |  | 0.9 |  | 0.9 |  |  |
| D5P | 12,447.00 |  | 22.06 |  | 97.8 |  | 2.3 |  | 1.5 |  | 196,148.00 |  | 20.09 |  | 0.5 |  | 0.7 |  | 0.6 |  |  |
| **D6P** | 6,443.00 |  | 12.05 |  | 94 |  | 0.8 |  | 0.6 |  | 129,620.00 |  | 13.25 |  | 0.3 |  | 0.4 |  | 0.4 |  | x |
| **D7P** | 2,238.00 |  | 3.75 |  | 83 |  | 2.4 |  | 2.5 |  | 36,097.00 |  | 3.58 |  | 1.4 |  | 2.1 |  | 2.4 |  | x |
| D8P | 14,498.00 |  | 25.26 |  | 97.9 |  | 2.4 |  | 1.9 |  | 213,204.00 |  | 21.61 |  | 0.6 |  | 0.7 |  | 0.7 |  |  |
| D9P | 10,805.00 |  | 18.96 |  | 98 |  | 2 |  | 1.6 |  | 163,064.00 |  | 16.71 |  | 0.5 |  | 0.8 |  | 1 |  |  |
| D10P | 12,327.00 |  | 22.16 |  | 97.2 |  | 1.9 |  | 1.3 |  | 213,402.00 |  | 21.8 |  | 0.5 |  | 0.5 |  | 0.5 |  |  |
| All P | 10,526.60 | 4,207.48 | 18.64 | 7.49 | 95.95 | 4.71 | 1.94 | 0.5 | 1.41 | 0.53 | 170,591.40 | 63,141.66 | 17.43 | 6.58 | 0.61 | 0.31 | 0.8 | 0.49 | 0.84 | 0.58 |  |
| D1NP | 11,766.00 |  | 20.02 |  | 98.3 |  | 2.3 |  | 1.6 |  | 221,420.00 |  | 22.82 |  | 0.5 |  | 0.7 |  | 0.8 |  |  |
| D2NP | 9,279.00 |  | 15.68 |  | 99.1 |  | 1.6 |  | 0.9 |  | 154,184.00 |  | 15.92 |  | 0.5 |  | 0.5 |  | 0.5 |  |  |
| D3NP | 9,033.00 |  | 15.6 |  | 98.4 |  | 2.5 |  | 1.6 |  | 155,290.00 |  | 15.84 |  | 0.8 |  | 0.9 |  | 0.9 |  |  |
| D4NP | 10,654.00 |  | 17.62 |  | 96.6 |  | 2.7 |  | 2.1 |  | 207,312.00 |  | 21.1 |  | 0.7 |  | 0.8 |  | 1 |  |  |
| D5NP | 7,539.00 |  | 12.76 |  | 98.7 |  | 2.7 |  | 2.4 |  | 157,185.00 |  | 16.46 |  | 0.6 |  | 0.8 |  | 0.9 |  |  |
| D6NP | 10,514.00 |  | 17.41 |  | 96.7 |  | 2.3 |  | 1.8 |  | 178,031.00 |  | 18.34 |  | 0.5 |  | 0.8 |  | 0.7 |  |  |
| D7NP | 8,878.00 |  | 14.07 |  | 98.7 |  | 2.1 |  | 1.8 |  | 165,344.00 |  | 16.87 |  | 0.7 |  | 0.7 |  | 0.8 |  |  |
| D8NP | 10,965.00 |  | 18.09 |  | 99.5 |  | 3.8 |  | 2.8 |  | 211,734.00 |  | 21.67 |  | 0.4 |  | 0.5 |  | 0.6 |  |  |
| D9NP | 7,387.00 |  | 12.28 |  | 98.2 |  | 3.1 |  | 1.6 |  | 133,188.00 |  | 13.5 |  | 0.8 |  | 1.1 |  | 1.1 |  |  |
| D10NP | 7,233.00 |  | 12.21 |  | 97.5 |  | 2.5 |  | 1.5 |  | 131,224.00 |  | 13.38 |  | 0.7 |  | 0.9 |  | 0.7 |  |  |
| All NP | 9,324.80 | 1,613.74 | 15.57 | 2.71 | 98.17 | 0.96 | 2.56 | 0.59 | 1.81 | 0.52 | 171,491.20 | 32,199.59 | 17.59 | 3.32 | 0.62 | 0.14 | 0.77 | 0.18 | 0.8 | 0.18 |  |
| DpoolP | 76,496.00 |  | 154.17 |  | 96 |  | 1.7 |  | 1.6 |  | 1,353,971.00 |  | 161.55 |  | 0.6 |  | 0.8 |  | 0.8 |  |  |
| DpoolNP | 71,022.00 |  | 141.52 |  | 97.5 |  | 2.2 |  | 1.5 |  | 1,246,168.00 |  | 146.63 |  | 0.6 |  | 0.8 |  | 0.9 |  |  |
| All pooled | 73,759.00 | 3,870.70 | 147.85 | 8.94 | 96.75 | 1.06 | 1.95 | 0.35 | 1.55 | 0.07 | 1,300,069.50 | 76,228.23 | 154.09 | 10.55 | 0.6 | 0 | 0.8 | 0 | 0.85 | 0.07 |  |

**Table S4:** Summary of variable and invariable regions.

|  | Library | Invariable |  | Variable |  |
| --- | --- | --- | --- | --- | --- |
|  |  | Regions | Proportion | Regions | Proportion |
| *Corbicula fluminea* | Individual | 10 | 0.001 | 7822 | 0.999 |
|  | Pooled | 26596 | 0.055 | 456310 | 0.945 |
|  | Computationally pooled | 18786 | 0.199 | 75448 | 0.801 |
| *Dreissena polymorpha* | Individual | 1 | 0.001 | 1149 | 0.999 |
|  | Pooled | 38717 | 0.048 | 760027 | 0.952 |
|  | Computationally pooled | 58231 | 0.305 | 132844 | 0.695 |

**Table S5:** Distribution of regions by mean difference in methylation. For each species and each of individual, pooled and computationally pooled data we indicate the number and proportion of regions according to the percentile bins for the mean difference in methylation between the polluted and non-polluted treatment groups.

| Species | Libraries | Quantile | MDM bin (%) | Power > 0.8 |  | Power < 0.8 |  |
| --- | --- | --- | --- | --- | --- | --- | --- |
|  |  |  |  | Regions | Proportion | Regions | Proportion |
| *C. fluminea* | Individual | 0 - 5% | [0.0387,0.0833) |  |  | 391 | 0.05 |
|  | Individual | 5 - 10% | [0.0833,0.212) |  |  | 1173 | 0.15 |
|  | Individual | 10 - 25% | [0.212,0.521) |  |  | 1955 | 0.25 |
|  | Individual | 25 - 50% | [0.521,1.39) |  |  | 1955 | 0.25 |
|  | Individual | 50 - 75% | [1.39,3.45) |  |  | 1173 | 0.15 |
|  | Individual | 75 - 90% | [3.45,5) |  |  | 391 | 0.05 |
|  | Individual | 90 - 95% | [5,18.4] |  |  | 392 | 0.05 |
|  | Individual | 95 - 100% | [9.33e-06,0.0387) |  |  | 392 | 0.05 |
|  | Pooled | 0 - 5% | [0.0837,0.163) |  |  | 22813 | 0.05 |
|  | Pooled | 5 - 10% | [0.163,0.422) | 33 | 0.00 | 68416 | 0.15 |
|  | Pooled | 10 - 25% | [0.422,1.15) | 3905 | 0.01 | 110172 | 0.24 |
|  | Pooled | 25 - 50% | [1.15,3.46) | 13011 | 0.03 | 101066 | 0.22 |
|  | Pooled | 50 - 75% | [11.1,88] | 9692 | 0.02 | 13124 | 0.03 |
|  | Pooled | 75 - 90% | [3.46,7.63) | 6691 | 0.01 | 61756 | 0.14 |
|  | Pooled | 90 - 95% | [7.63,11.1) | 4005 | 0.01 | 18810 | 0.04 |
|  | Pooled | 95 - 100% | [9.39e-06,0.0837) |  |  | 22816 | 0.05 |
|  | Computationally pooled | 0 - 5% | [0.121,0.223) |  |  | 3772 | 0.05 |
|  | Computationally pooled | 5 - 10% | [0.223,0.529) | 23 | 0.00 | 11294 | 0.15 |
|  | Computationally pooled | 10 - 25% | [0.529,1.47) | 492 | 0.01 | 18346 | 0.24 |
|  | Computationally pooled | 25 - 50% | [1.47,4.51) | 1448 | 0.02 | 17438 | 0.23 |
|  | Computationally pooled | 50 - 75% | [10.7,16) | 947 | 0.01 | 2825 | 0.04 |
|  | Computationally pooled | 75 - 90% | [16,100] | 1945 | 0.03 | 1828 | 0.02 |
|  | Computationally pooled | 90 - 95% | [4.51,10.7) | 1611 | 0.02 | 9706 | 0.13 |
|  | Computationally pooled | 95 - 100% | [9.33e-06,0.121) |  |  | 3773 | 0.05 |
| *D. polymorpha* | Individual | 0 - 5% | [0.000541,0.0363) |  |  | 58 | 0.05 |
|  | Individual | 5 - 10% | [0.0363,0.0647) |  |  | 57 | 0.05 |
|  | Individual | 10 - 25% | [0.0647,0.168) |  |  | 172 | 0.15 |
|  | Individual | 25 - 50% | [0.168,0.407) |  |  | 287 | 0.25 |
|  | Individual | 50 - 75% | [0.407,1.38) |  |  | 287 | 0.25 |
|  | Individual | 75 - 90% | [1.38,5.01) |  |  | 173 | 0.15 |
|  | Individual | 90 - 95% | [5.01,8.09) |  |  | 57 | 0.05 |
|  | Individual | 95 - 100% | [8.09,39.1] |  |  | 58 | 0.05 |
|  | Pooled | 0 - 5% | [0.0798,0.154) |  |  | 38000 | 0.05 |
|  | Pooled | 5 - 10% | [0.154,0.409) | 110 | 0.00 | 113895 | 0.15 |
|  | Pooled | 10 - 25% | [0.409,1.21) | 12640 | 0.02 | 177366 | 0.23 |
|  | Pooled | 25 - 50% | [1.21,4.4) | 27941 | 0.04 | 162056 | 0.21 |
|  | Pooled | 50 - 75% | [12.6,87.9] | 26269 | 0.03 | 11733 | 0.02 |
|  | Pooled | 75 - 90% | [2.01e-05,0.0798) |  |  | 38002 | 0.05 |
|  | Pooled | 90 - 95% | [4.4,8.91) | 29197 | 0.04 | 84817 | 0.11 |
|  | Pooled | 95 - 100% | [8.91,12.6) | 17617 | 0.02 | 20384 | 0.03 |
|  | Computationally pooled | 0 - 5% | [0.000324,0.19) | 1 | 0.00 | 6613 | 0.05 |
|  | Computationally pooled | 5 - 10% | [0.19,0.325) | 13 | 0.00 | 6617 | 0.05 |
|  | Computationally pooled | 10 - 25% | [0.325,0.78) | 153 | 0.00 | 19783 | 0.15 |
|  | Computationally pooled | 25 - 50% | [0.78,2.36) | 1067 | 0.01 | 32142 | 0.24 |
|  | Computationally pooled | 50 - 75% | [16.2,23.9) | 4052 | 0.03 | 2590 | 0.02 |
|  | Computationally pooled | 75 - 90% | [2.36,7.43) | 3701 | 0.03 | 29512 | 0.22 |
|  | Computationally pooled | 90 - 95% | [23.9,100] | 6048 | 0.05 | 595 | 0.00 |
|  | Computationally pooled | 95 - 100% | [7.43,16.2) | 6647 | 0.05 | 13279 | 0.10 |

## Supplemental Figures

**Figure S1:**
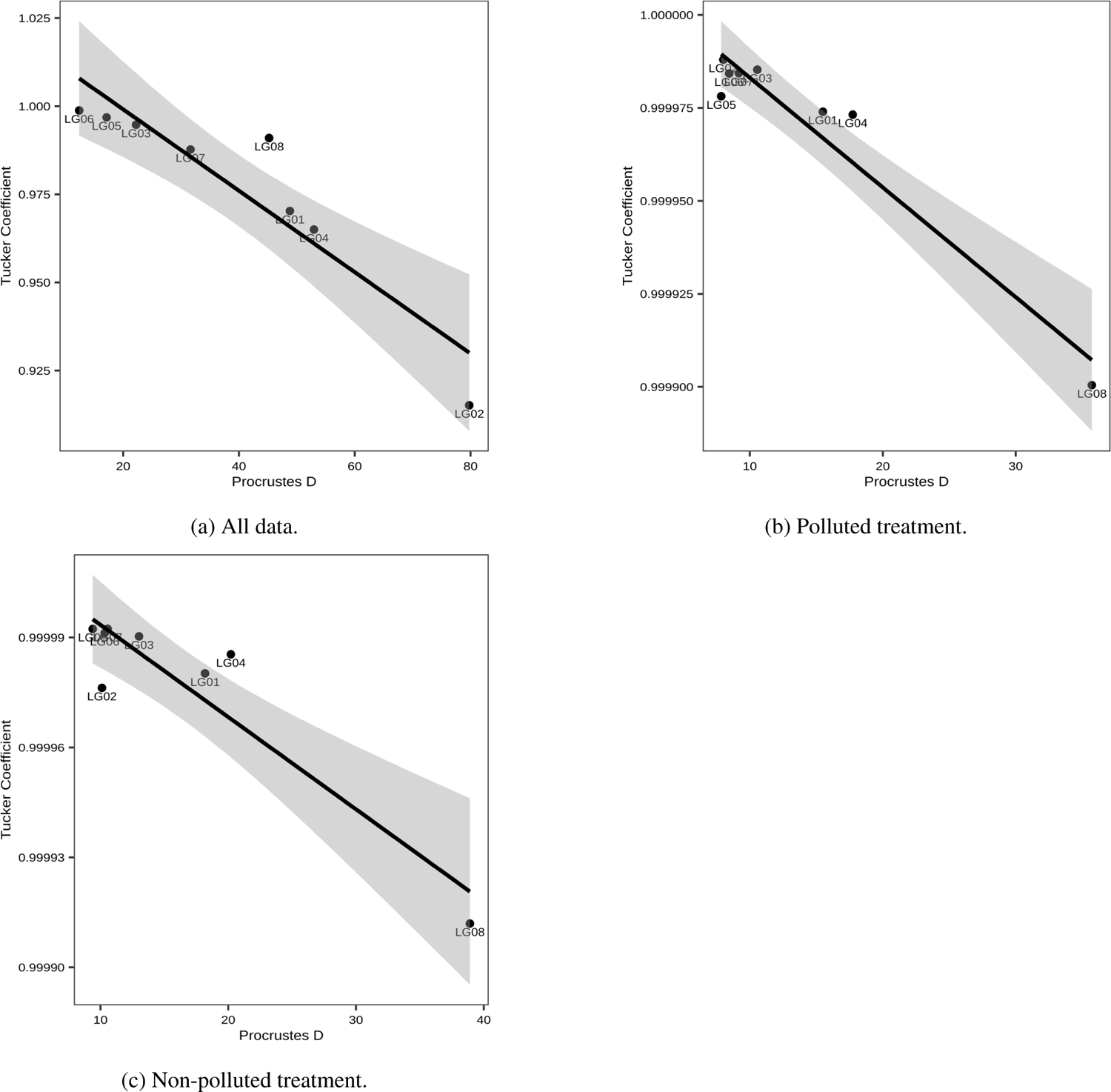
Correlation of the Tucker’s coefficient and the Procrustes transformation D for *C. fluminea* using samples from the *in vitro* pooled and individual libraries with tile-resolution data. Data subsets indicated for each plot.

**Figure S2:**
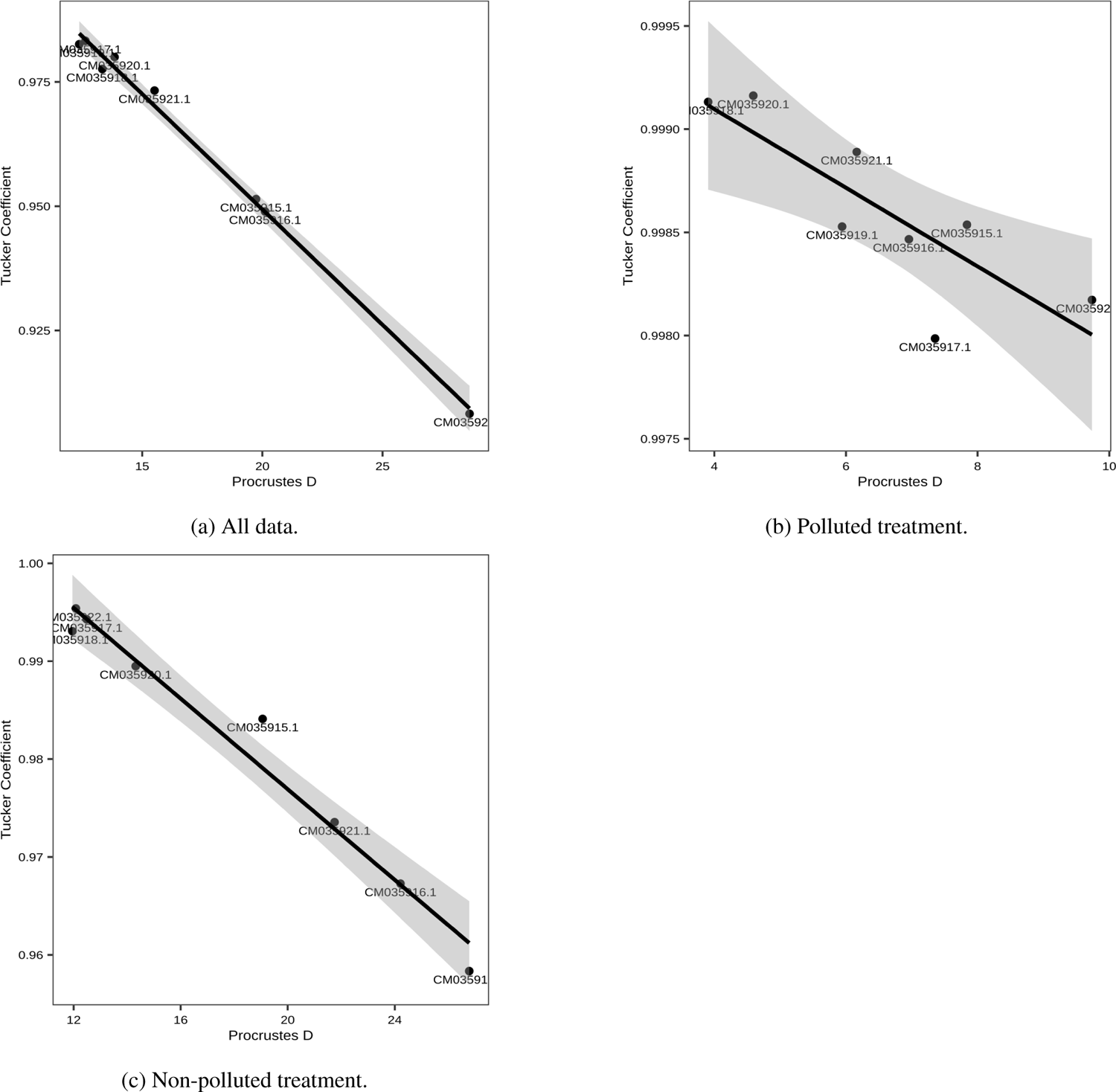
Correlation of the Tucker’s coefficient and the Procrustes transformation D for *D. polymorpha* using samples from the *in vitro* pooled and individual libraries with tile-resolution data. Data subsets indicated for each plot.

## Notes

### Competing Interest Statement

The authors have declared no competing interest.

